# MxtR/ErdR is a central regulator of short-chain fatty acid metabolism in *Pseudomonas alloputida*

**DOI:** 10.64898/2026.09.07.749794

**Authors:** Fabienne Burr, Tania Henríquez, Michelle Eder, Ivan Cerkez, Siobhan A. Cusack, Isabella Heinzelmann, Heinrich Jung

**Author notes:** Corresponding author: Heinrich Jung, Ludwig-Maximilians-Universität München, Fakultät für Biologie, Mikrobiologie, Biozentrum, Großhaderner Straße 2-4, 82152 Planegg-Martinsried, Germany. Università degli Studi di Siena, Siena, Italy (Tania Henríquez).

## Abstract

Short-chain fatty acids (SCFAs) such as acetate and propionate represent important carbon and energy sources for environmental pseudomonads, requiring coordinated regulation to ensure efficient assimilation while maintaining metabolic homeostasis. In *Pseudomonas alloputida* KT2440, the two-component system MxtR/ErdR (CrbS/CrbR) is known to activate acetate assimilation through regulation of *acsA-I*. However, the full extent of the MxtR/ErdR regulon and its physiological role beyond acetate metabolism have remained unclear. We demonstrate that MxtR/ErdR directly regulates the methylcitrate cycle and coordinates both acetate and propionate metabolism. To define the MxtR/ErdR regulatory network, we combined comparative transcriptomics, physiological analyses, promoter-reporter assays, electrophoretic mobility shift assays (EMSAs) and targeted mutagenesis. Comparative transcriptomic analyses of the *mxtR*-H806N and Δ*erdR* mutants revealed extensive changes in gene expression, including coordinated downregulation of genes involved in propionate metabolism alongside genes associated with central carbon metabolism, transport, chemotaxis, and signal transduction. Consistent with these transcriptional changes, deletion of either *mxtR* or *erdR* abolished growth on propionate. Promoter-reporter assays and EMSAs demonstrated direct binding of ErdR to a conserved imperfect inverted repeat upstream of the *prp* gene cluster and *prpE*. Mutational analyses confirmed the functional importance of this binding motif for promoter activation. In addition, MxtR/ErdR contributed to pyruvate utilization through regulation of transport-associated genes, whereas deletion of downstream target genes only caused modest phenotypes, indicating that the physiological effects of MxtR/ErdR arise from coordinated regulation of multiple pathways. Our findings substantially expand the MxtR/ErdR regulon and identify this signaling system as a central regulator of SCFA homeostasis in *P. alloputida* KT2440.

## 1. Introduction

Short-chain fatty acids (SCFAs), including acetate, propionate, and butyrate, are key intermediates of microbial carbon metabolism. Their assimilation provides bacteria with substantial metabolic flexibility, supporting persistence and competitive growth under fluctuating nutrient conditions in diverse environments, including soil, aquatic ecosystems, and plant- and animal-associated habitats (Wolfe, 2005; Koh et al., 2016). However, elevated SCFA concentrations can be detrimental to bacterial cells. In their undissociated form, SCFAs can diffuse across the cytoplasmic membrane and dissociate in the cytoplasm, thereby dissipating the proton motive force and causing intracellular anion accumulation (Russell and Diez-Gonzalez, 1998; Sun and O’Riordan, 2013; Pinhal et al., 2019). Moreover, activation of SCFAs by CoA ligation alters the intracellular acyl-CoA/CoA-SH balance, potentially limiting CoA availability for other essential metabolic reactions. Accumulation of metabolic intermediates, such as acetyl phosphate, further affects cellular physiology by modulating protein acetylation and enzyme activity (Wolfe, 2005; Pinhal et al., 2019; Ren et al., 2019). Consequently, SCFA uptake and metabolism must be tightly regulated to maximize their nutritional benefits while preventing metabolic imbalance and toxicity.

Acetate utilization is controlled by the so-called acetate switch, which marks the transition from acetate excretion to acetate assimilation. This transition requires induction and activation of the acetate assimilation machinery (Wolfe, 2005). In *Escherichia coli*, the acetate switch is primarily mediated through transcriptional activation of *acs*, encoding acetyl-CoA synthetase, by the global regulators CRP and FNR in response to carbon availability and oxygen limitation (Kumari et al., 2000). Acetyl-CoA synthetase activity is additionally regulated by reversible lysine acetylation (Wolfe, 2005; Castaño-Cerezo et al., 2015), while acetate assimilation is further coordinated through regulation of downstream pathways, including the glyoxylate shunt via IclR and FadR (Kumari et al., 2000).

In other Gammaproteobacteria, the acetate switch is controlled by the two-component system (TCS) CrbS/CrbR, designated MxtR/ErdR in *Pseudomonas* species, composed of the membrane-bound sensor kinase MxtR and its cytosolic response regulator ErdR. This signaling system activates transcription of *acs* homologues and is therefore essential for acetate assimilation. In *Vibrio cholerae*, CrbS/CrbR-dependent acetate utilization influences host metabolism in *Drosophila melanogaster* and contributes to bacterial virulence (Hang et al., 2014). A conserved function has also been demonstrated in several *Pseudomonas* species, including *Pseudomonas aeruginosa*, *P. entomophila*, *P. fluorescens*, and *P. alloputida*, where MxtR/ErdR controls expression of *acsA* and is required for growth on acetate as the sole carbon source (Jacob et al., 2017; Sepulveda and Lupas, 2017; Henriquez and Jung, 2021). Recently, the orthologous TCS AcmS/AcmR was shown to regulate acetate metabolism in *Acinetobacter baumannii* (Pokhrel et al., 2026). Comparative promoter analyses identified a conserved imperfect inverted repeat that is required for MxtR/ErdR-dependent *acs* expression and is presumed to represent the ErdR binding site. Based on this sequence motif, several additional genes were proposed as potential ErdR targets, although experimental evidence supporting their regulation has remained limited (Sepulveda and Lupas, 2017).

We use the rhizosphere bacterium and industrial chassis *P. alloputida* KT2440 (Weimer et al., 2020; Martinez-Garcia and de Lorenzo, 2024) as a model organism. Our previous targeted gene expression analyses and electrophoretic mobility shift assays demonstrated that ErdR controls the expressions of *acsA-I*, *actP-I*, *scpC*, and PP_0354 (Henriquez and Jung, 2021; Henriquez et al., 2021; Henriquez et al., 2023). As expected, *acsA-I* is essential for acetate utilization. In contrast, deletion of *actP-I* has little effect on growth at millimolar acetate concentrations, but impairs growth on pyruvate, which depends on ActP-I for efficient uptake. ScpC, an acetate:succinate CoA transferase, catalyzes CoA transfer from succinyl-CoA to acetate and is proposed to substitute for succinyl-CoA synthetase in the tricarboxylic acid (TCA) cycle under elevated acetate concentration conditions, thereby contributing to acetate tolerance (Henriquez et al., 2023). These findings indicate that MxtR/ErdR regulates a broader physiological response than acetate assimilation alone.

In the present study, we sought to define the MxtR/ErdR regulon and to clarify its physiological significance in *P. alloputida* KT2440. Comparative transcriptome analyses of wild-type cells and *mxtR*-H806N and Δ*erdR* mutants identified numerous differentially expressed genes, including those involved in central carbon metabolism, signal transduction, chemotaxis and membrane transport. Notably, several MxtR/ErdR-dependent genes are associated with propionate utilization. Consistent with these transcriptional changes, both *mxtR* and *erdR* proved essential for growth on propionate. We further demonstrate that expression of the methylcitrate cycle (2-MCC) genes, which mediate degradation of toxic propionyl-CoA generated during the metabolism of odd-chain fatty acids, depends on MxtR/ErdR. Electrophoretic mobility shift assays indicate direct binding of ErdR to the promoter regions of both the 2-MCC operon and *prpE*, encoding a propionyl-CoA synthetase. Together, our results substantially expand the known MxtR/ErdR regulatory network and reveal that this signaling system coordinates not only acetate assimilation, but also propionate metabolism in *P. alloputida* KT2440.

## 2. Material & Methods

### 2.1. Bacterial strains and cultivation conditions

All bacterial strains and plasmids used in the scope of this study are listed in Tables S1 and S2. *Pseudomonas alloputida* strains were cultivated under aerobic conditions, consistent shaking at 180 rpm, at 30°C in either King’s B (KB) medium (King et al., 1954) containing 2% peptone, 1% (w/v) glycerol, 0.15% K_2_HPO_4_, and 0.15% MgSO_4_x7H_2_O, or M9 minimal medium as previously described by Henriquez et al (2023) supplemented with 20 or 40 mM carbon source (acetate, propionate, succinate, pyruvate, or glucose). Cells transformed with plasmids for complementation (pUCP20-ANT2-MCS (Hoffmann et al., 2021), pSEVA224 (Silva-Rocha et al., 2013), cloning purposes (pNPTS138-R6KT (Lassak et al., 2010)), gene expression analysis (pBBR1-MCS5-*lux* (Gödeke et al., 2011)) or protein overexpression (pET16b) were supplemented with 50 μg/ml kanamycin (pUCP20-ANT2-MCS, pSEVA224), 30 μg/ml gentamicin (pBBR1-MCS5-*lux*) or 100 µg/ml ampicillin. Expression from the plasmids was induced with 0.5 mM isopropyl β-D-1-thiogalactopyranoside (IPTG; pSEVA224, pET16b) or 5 µM anthranilic acid (2AA; pUCP20).

### 2.2. Generation of plasmids and mutants

Alterations of the genome (gene deletions, nucleotide substitutions) were generated via double homologous recombination with the pNPTS138-R6KT suicide plasmid, as previously described (Lassak et al., 2010; Henriquez et al., 2019). All oligonucleotides used for this and further purposes are listed in Table S3. For complementation purposes, the gene of interest was amplified via polymerase chain reaction (PCR) and cloned into the multiple cloning site (MCS) of pUCP20-ANT2-MCS or pSEVA224. For gene expression analyses, promoter regions were amplified and cloned into pBBR1-MCS5-*lux*. All plasmids and deletion strains were verified via sequencing.

### 2.3. Cultivation and RNA isolation for RNA sequencing

*Pseudomonas alloputida* strains (KT2440 (wild type), *mxtR*-H806N, Δ*erdR*) were pre-cultured aerobically in King’s B medium overnight. Cells were then transferred to a baffled flask for an over-day culture in M9 minimal medium with 20 mM succinate, with an initial OD_600_ of 0.05. Cells were grown aerobically, under continuous shaking (180 rpm) at 30°C until an OD_600_ of 0.5 was reached. Cells were then centrifuged at 5000 rpm, 30°C for 60 seconds and shifted to M9 minimal medium with 40 mM acetate. Cells were further incubated for 15 min under continuous shaking (180 rpm) at 30°C. Then, bacterial strains were mixed with a solution of phenol (1% v/v final concentration) and ethanol (20% v/v final concentration) and shock-frozen in liquid nitrogen. Cells were stored at -80°C until RNA isolation occurred. RNA isolation followed the phenol-chloroform-isoamyl alcohol (PCI) protocol (Sambrook and Russell, 2006), with some modifications (Petrov et al., 2022), as previously described (Riquelme-Barrios et al., 2025).

Extracted RNA concentrations were determined using the Nanodrop 1000 spectrophotometer (Thermo Fisher Scientific, Waltham, MA, USA). RNA integrity and quality were verified via chip gel electrophoresis using a 2100 Bioanalyzer and an RNA Nano Chip Kit (Agilent Technologies, Santa Clara, CA, USA). The extracted RNA was then subjected to a DNase I treatment, followed by an RNA Clean-up and subsequent quality testing with the 2100 Bioanalyzer. RNA concentrations were determined using the Qubit 4 Fluorometer and the corresponding Qubit RNA High Sensitivity Assay Kit (Thermo Fisher Scientific, Waltham, MA, USA). RNA was diluted to 20 ng/µl and sequenced by Novogene Sequencing Germany (Planegg, Germany) using the paired-end Prokaryotic mRNA seq protocol to produce a minimum of 10 million reads.

### 2.4. Downregulated gene functional characterization

Differential gene expression values in the mutant lines compared to the wild type were calculated by Novogene Sequencing Germany (Planegg, Germany). Genes were filtered to retain those that were significantly differentially expressed at an adjusted *p*-value <0.05. Then the 50 most strongly downregulated genes in each mutant (as determined using log_2_(fold change) values compared to the wild type) were selected for visualization. Treemaps were generated with Proteomaps (Liebermeister et al., 2014) using annotations from the user-defined template “*Pseudomonas putida* KT2440 KO (adapted by N 12072024)”. For genes of interest that were not present in this file, annotations were obtained from the KEGG BRITE Database: KEGG Orthology (BRITE identifier: ppu00001) (Kanehisa, 2017; Laboratories, 2026); genes of interest also without annotations in the KEGG Orthology file were assigned functions based on a manual literature search or given the designation “Function unknown”. The colors and text of the graphs produced by Bionic Vis were manually adjusted in Adobe Illustrator for contrast and visibility.

### 2.5. Growth analyses

To investigate growth, overnight cultures of the strain of interest were prepared in KB medium. If the strains carried a plasmid, the appropriate antibiotic was added. Over-day cultures in M9 minimal medium with 20 mM succinate were inoculated with an initial OD_600_ of 0.4, using the overnight cultures. This was performed in order to ensure that cells used for the growth analyses were in the exponential growth phase. M9 minimal medium containing a single carbon source, as well as the appropriate antibiotic, was inoculated with the pre-cultured cells in 96-well plates with a starting OD_600_ of 0.1. Cells were incubated at 30°C with double orbital shaking at 600 rpm in a CLARIOstar Plus plate-reader (BMG LABTECH, Ortenberg, Germany). Growth was determined by measuring OD_600_ hourly. Growth curves, area under the curve (AUC) and statistical analyses were generated and calculated using GraphPad Prism version 10.4.1 for Windows, GraphPad Software, Boston, MA, USA.

### 2.6. qRT-PCR

Cell cultivation, RNA isolation and preparation were performed as described above for RNA sequencing. RNA quantification was performed with the Nanodrop 1000 spectrophotometer (Thermo Fisher Scientific, Waltham, MA, USA). Reverse transcription of 0.5 µg of RNA was performed with the High-Capacity cDNA Reverse Transcription Kit (Thermo Fisher Scientific, Waltham, MA, USA). cDNA was diluted 1:5 using nuclease-free water in order to be used as samples for qPCR. The qPCR reaction mixture contained diluted cDNA, corresponding primers (see Table S3), as well as iQ SYBR Green Supermix (Bio-Rad Laboratories, Hercules, CA, USA). The reaction was performed in a C1000 Touch Thermal Cycler (Bio-Rad Laboratories, Hercules, CA, USA). C_t_ values were normalized using the reference gene *gapA* (glyceraldehyde 3-phosphate dehydrogenase).

### 2.7. Luciferase activity assay

Overnight cultures of the *P. alloputida* strains of interest (KT2440 (wild type), Δ*mxtR*, and Δ*erdR*) carrying either pBBR1-P_PP_2333_-*lux*, pBBR1-P_PP_2333-IR-[tga]_-*lux,* pBBR1-P_PP_2333-IR-[del]_-*lux,* pBBR1-P*_prpE_-lux,* pBBR1-P*_prpE_*_-IR-[tct]_-*lux* or pBBR1-MCS5*-lux,* cultivated in KB (with 30 µg/ml gentamicin), were used to inoculate over day cultures in M9 minimal medium with 20 mM succinate at 30°C (start OD_600_ of 0.4). Ninety-six-well plates containing M9 minimal medium supplemented with either 20 mM propionate or 20 mM succinate as a sole carbon source, as well as 30 µg/ml gentamicin and 0.5 mM IPTG were inoculated with a start OD_600_ of 0.1. Growth and luminescence were measured quarter-hourly in the CLARIOstar Plus (BMG LABTECH, Ortenberg, Germany) plate-reader. Cells were incubated at 30°C with double orbital shaking, 600 rpm. Growth curves, RLU/OD_600_ curves, area under the curve (AUC) and statistical analyses were generated and calculated using GraphPad Prism version 10.4.1 for Windows, GraphPad Software, Boston, MA, USA.

### 2.8. ErdR purification

The gene *erdR* (PP_1635) was cloned into the pET16b vector, introducing an N-terminal polyhistidine tag (10xHis). *E. coli* C41 cells were transformed with the pET16b-*erdR* vector. Three liters of LB medium (with 100 µg/ml of carbenicillin) were inoculated with 20 ml of an overnight culture of C41 pET16b-*erdR* per liter of medium and incubated at 30°C, 180 rpm until an OD_600_ between 0.55 and 0.6 was reached. Overexpression of 10H-ErdR was then induced with 0.5 mM IPTG (3 h, 30 °C, 180 rpm). After induction, cells were harvested at 6500 rpm, 15 min, 4°C and pellets were washed with 50 mM Tris/HCl (pH 8). Cell pellets were flash-frozen and stored at -80°C until further use.

Cell pellets were thawed on ice, resuspended in the appropriate buffer (50 mM Tris/HCl pH 8, 300 mM KCl, 10 mM imidazole pH 8, 0.5 mM phenylmethylsulfonyl fluoride, DNase I) and disrupted in a constant cell disrupter at 1.35 kbar in two runs. Cell debris was removed via centrifugation at 4,500 x g, 4°C, 1h. Cytosol was separated from the membrane fraction via ultra-centrifugation at 184,000 x g, 4°C, 1 h.

For initial purification via affinity chromatography, Ni-nitrilotriacetic acid (Ni-NTA) resin was transferred to a gravity flow chromatography column and equilibrated with 50 mM Tris/HCl pH 8, 300 mM KCl, 10 mM imidazole pH 8. Afterwards, the cytosol was loaded onto the column. In four washing steps, imidazole concentrations of the washing buffer (50 mM Tris/HCl pH 8, 5% glycerol, 300 mM KCl) were gradually increased in 20-mM steps, from an initial concentration of 10 mM up to 70 mM, to remove unspecific binding of non-target proteins. 10H-ErdR was eluted after 30 min of incubation with the elution buffer (50 mM Tris/HCl pH 8, 5% glycerol, 300 mM KCl, 200 mM imidazole pH 8). The concentration of protein in the eluted fractions was determined via Bradford protein assay (Bradford, 1976).

To increase purity of the protein eluate and remove imidazole residues, affinity chromatography was followed up with size exclusion chromatography. Thus, the Superdex 200 Increase 10/300 GL column (Cytiva, Marlborough, MA, USA) was equilibrated with 50 mM Tris/HCl pH 8, 5% glycerol and 300 mM KCl using the ÄKTA pure chromatography system (Cytiva, Marlborough, MA, USA). Before loading the Ni-NTA elution fraction, it was centrifuged for 10 min, 4°C at 21,130 x g to pellet any aggregates. Five hundred µl of 10H-ErdR were loaded onto the column and eluted in 500 µl fractions at 0.3 ml/min (elution buffer 50 mM Tris/HCl pH 8, 5% glycerol and 300 mM KCl). The Bradford protein assay (Bradford, 1976) was used to determine the protein concentrations of peak fractions.

Protein purity was assessed using a Coomassie-stained SDS-gel. The presence and identity of 10-ErdR was confirmed via western blot. For this purpose, the 0.2 µm nitrocellulose membrane was incubated with a mouse anti-6xHis primary antibody (Thermo Fisher Scientific, Waltham, MA, USA), followed by a chicken anti-mouse IgG (H+L) HRP secondary antibody (Thermo Fisher Scientific, Waltham, MA, USA). Bands were developed using the Clarity Western ECL Substrates (Bio-Rad Laboratories, Hercules, CA, USA) according to the manufacturer’s instructions. Blots were imaged using the Bio-Rad ChemiDoc Imaging System.

### 2.9. Electrophoretic Mobility Shift Assay (EMSA)

Promoter regions (around 120-160 bp upstream of start codon) of PP_2333, *prpE* and *scpC* were amplified via PCR using 5’-Cy5-labelled primers (Table S3). The promoter region of *scpC* was used as a positive control and an internal region of the superoxide dismutase *sodB* was used as a negative control. Binding and non-binding of ErdR to the positive and negative controls, respectively, has already been shown (Henriquez and Jung, 2021).

In each EMSA reaction, 1 nM of promoter region was added to variable concentrations of purified 10H-ErdR (0, 1, 3 and 5 µM). Prior to mixing of DNA with protein, 10H-ErdR was phosphorylated with 2 mM acetyl-phosphate for 30 min at room temperature, with the addition of EMSA buffer (1 mM dithiothreitol, 2 mM MgCl_2_, 10% glycerol, 10 mM Tris/HCl pH 8, 100 mM KCl, 0.05 mg/ml bovine serum albumin and 40 ng/µl sheared salmon sperm DNA). Promoter DNA and protein were mixed and incubated for 20 min at room temperature. The samples (10 µl) were loaded onto a 10% tris-glycine gel (NuSep, Germantown, MD, USA), which was pre-run at 80V in 1x tris-glycine buffer (250 mM Tris, 192 mM glycine, pH 8) for 1h. Gel electrophoresis of the samples was performed at 80V for 90 min. Gels were imaged using the Bio-Rad ChemiDoc Imaging System.

### 2.10. Statistical analyses

Statistical tests (ANOVA and Dunnett’s multiple comparison, as well as t-test) were performed using GraphPad Prism version 10.4.1 for Windows, GraphPad Software, Boston, MA, USA.

## 3. Results

### 3.1. Impacts of *mxtR* and *erdR* on the transcriptome of *P. alloputida* KT2440

To obtain a comprehensive overview of genes directly or indirectly regulated by the MxtR/ErdR TCS, the transcriptomes of *mxtR*-H806N and Δ*erdR* mutants were compared to that of wild type *P. alloputida* KT2440. The *mxtR*-H806N mutant, carrying a missense mutation in the histidine kinase A phosphoacceptor domain, represents an inactive *mxtR* mutant. All strains were cultivated on succinate and subsequently shifted to acetate for 15 min prior to RNA isolation.

Compared with the wild type, the *mxtR*-H806N mutant exhibited significant differential expression of 347 genes, including 164 downregulated (log₂ fold change <-1) and 183 upregulated genes (log₂ fold change >1, *p_adj_* < 0.05) (Fig. 1A). The Δ*erdR* mutant showed a smaller transcriptional response compared to the wild type, with 52 downregulated genes and six upregulated genes (Fig. 1B). Notably, 34 of the 50 most strongly downregulated genes were shared between the two mutants. KEGG pathway analysis revealed that the downregulated genes encode proteins involved in central carbon metabolism, including glycolysis/gluconeogenesis, fatty acid degradation, the TCA cycle and the glyoxylate shunt, as well as transport systems, signal transduction proteins and additional TCSs. However, the largest functional category comprised genes encoding proteins of unknown function (Fig. 1C-D).

**Fig. 1:**
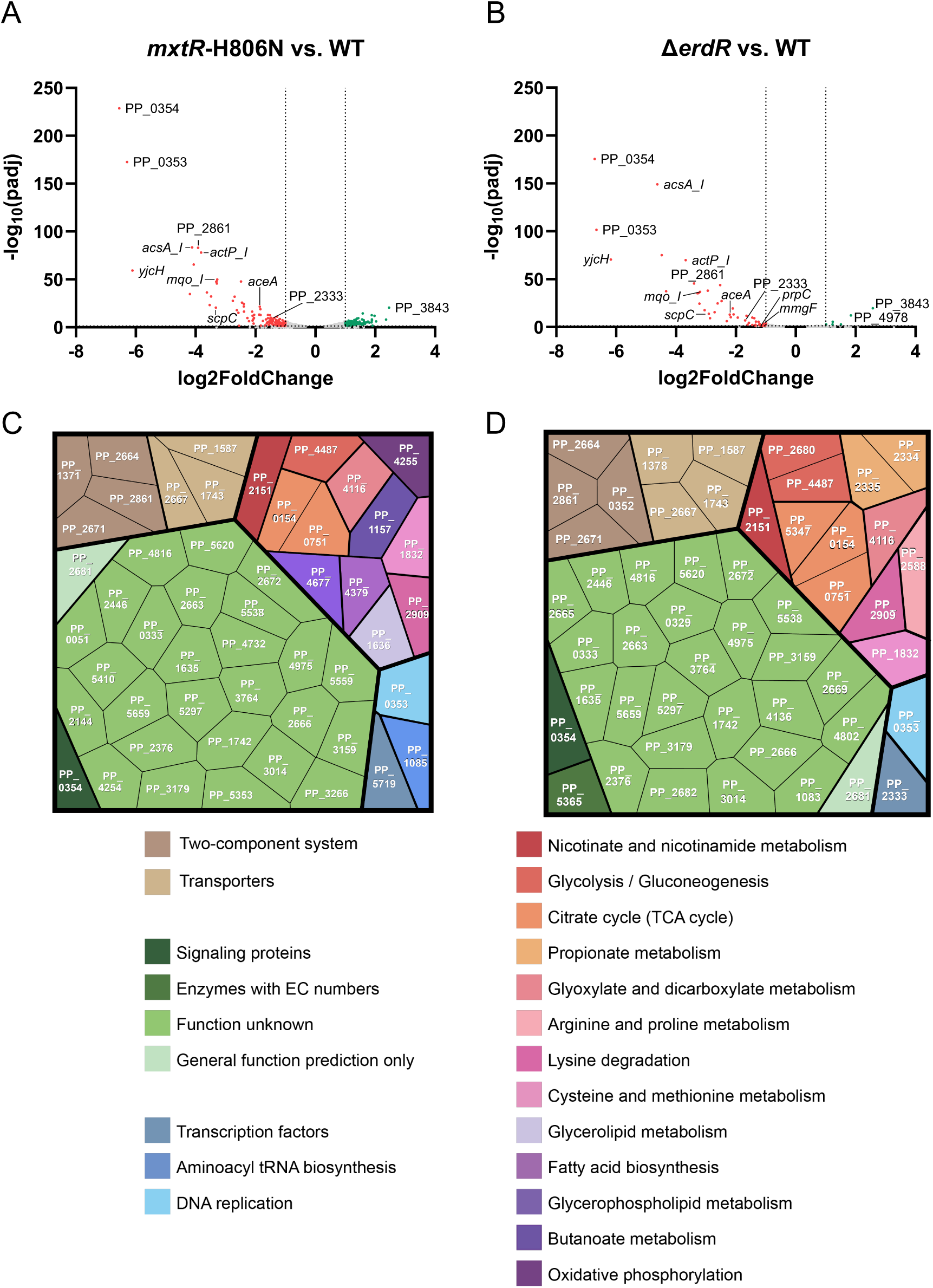
Gene expression profile of *mxtR*-H806N and Δ*erdR* compared to wild type *P. alloputida* KT2440. (A, B) Differential gene expression of *mxtR*-H806N and Δ*erdR* compared to wild type (WT). Cells were pre-cultured in M9 minimal medium with 20 mM succinate and shifted to M9 with 40 mM acetate for 15 min. Cultures were shock-frozen and RNA isolation followed promptly. Genes of interest investigated in this study are annotated. (C, D) Functional annotations of the 50 most strongly downregulated genes in (C) *mxtR*-H806N and (D) Δ*erdR* compared to the wild type. Downregulated genes that have been functionally annotated were generally associated with central carbon metabolism, including propionate metabolism in the Δ*erdR* mutant.

All previously known direct target genes of the TCS were consistently among the most strongly downregulated genes in both mutants, including the *yjcH - actP-I* operon (PP_1742-PP_1743), *acsA-I* (PP_4487) and *scpC* (PP_0154), which are involved in acetate transport and metabolism, as well as the PP_0353 - PP_0354 operon, which encodes putative regulators of acetate metabolism. In addition, several previously unrecognized genes were found among the most strongly downregulated genes (Table S4 and S6). These included PP_2861, encoding a chemoreceptor for C2 and C3 carboxylic acids (Garcia et al., 2015); *mqo-I* (PP_0751), encoding the TCA cycle enzyme malate:quinone oxidoreductase, which transfers electrons directly to the quinone pool of the respiratory chain, thereby bypassing complex I (Mellgren et al., 2009); *aceA* (PP_4116) encoding isocitrate lyase of the glyoxylate shunt; *pqqD-II* (PP_2681), required for the biosynthesis of the alcohol dehydrogenase cofactor pyrroloquinoline quinone (PQQ); and several genes of the *ped* cluster (PP_2663, PP_2666, PP_2667, and PP_2669), which are involved in the oxidation of a broad range of alcohols and aldehydes (Puiggene and Nikel, 2026). In addition, genes involved in propionate metabolism (PP_2333, *mmgF*, *prpC* and *prpE*) were significantly, although less strongly, downregulated in both mutants. Several downregulated genes also encoded putative signal transduction proteins, including the two-component systems PedS1/PedR1 (PP_2664/PP_2665) and PedS2/PedR2 (PP_2671/PP_2672).

In contrast to the extensive downregulation observed in both mutants, the observed changes for upregulated genes were generally modest (Table S5 and S7). These genes included several encoding proteins of unknown function (e.g., PP_3843 and PP_4978), components of the cellular stress response such as *groL* and *groS*, and genes that may functionally compensate for downregulated metabolic enzymes, including *mqo-II*. The list of the top 50 (if applicable) differentially expressed genes for both mutants, including their annotated functions, is provided in Tables S4 - S7.

### 3.2. MxtR/ErdR is required for growth of *P. alloputida* KT2440 on propionate

In light of the transcriptomic analysis revealing reduced expression of genes involved in propionate metabolism, we next focused on the role of MxtR/ErdR in consuming the C3 compound propionate. We first examined the growth of *P. alloputida* KT2440 and the derived mutants Δ*mxtR* and Δ*erdR* on different carbon sources. We found that individual deletion of *mxtR* or *erdR* stunted bacterial growth on 20 mM propionate as the sole carbon source (Fig. 2A), comparable to growth on minimal medium with acetate as a sole carbon source (Henriquez and Jung, 2021). Deletion of *mxtR* or *erdR* prevented growth on higher propionate concentrations, i.e., 40 mM propionate (Fig. 2B).

**Fig. 2:**
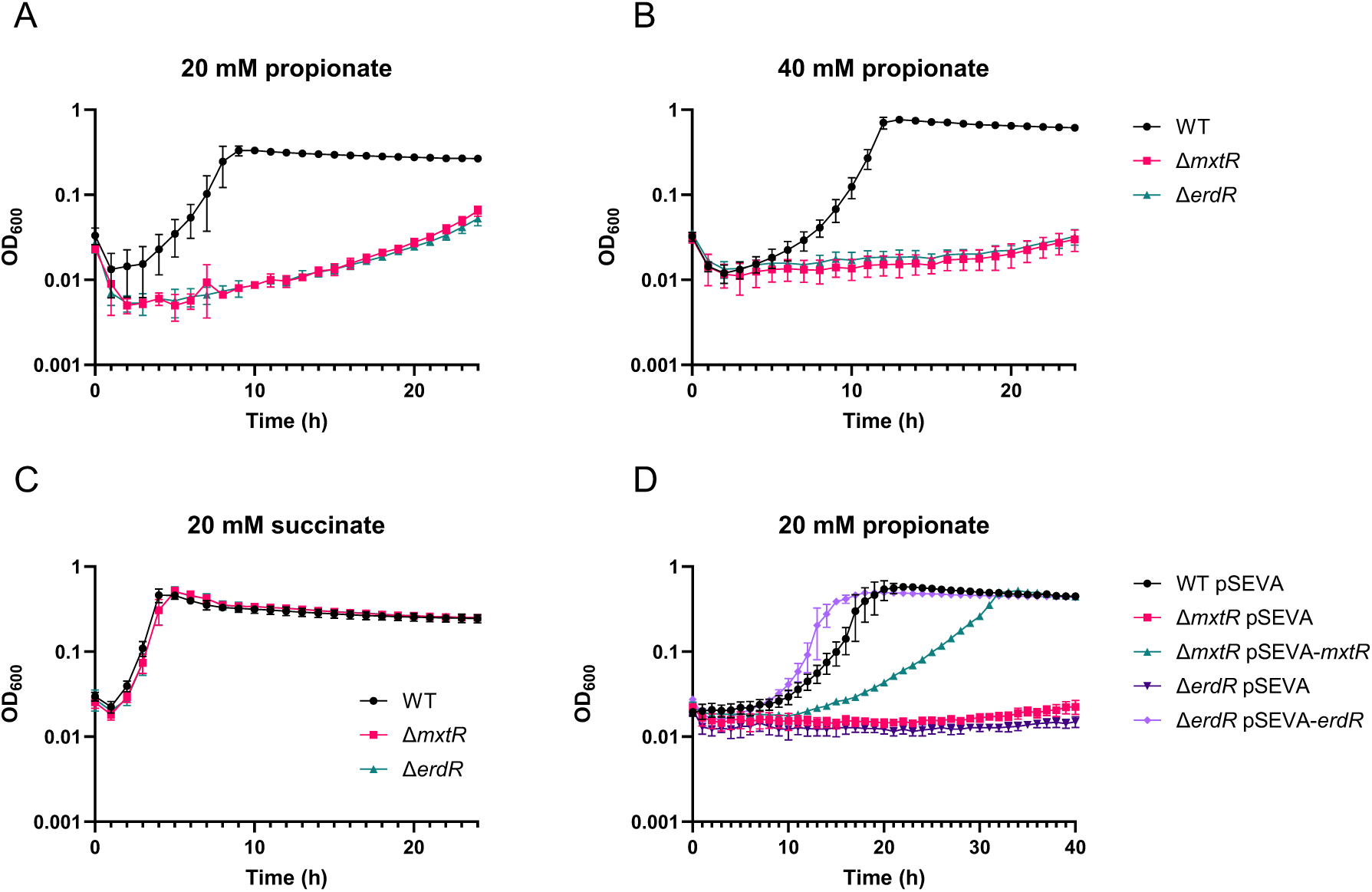
Impact of *mxtR* and *erdR* on growth in M9 minimal medium. (A) Time courses showing growth of wild type (WT), Δ*mxtR*, Δ*erdR* in M9 minimal medium supplemented with 20 mM propionate, (B) 40 mM propionate, and (C) 20 mM succinate. (D) Complementation of the growth defect on 20 mM propionate. Growth analyses were performed with an initial OD_600_ of 0.1 and cells were cultivated in 96-well plates in a CLARIOstar Plus plate-reader for 24 or 40 h, measuring OD_600_ hourly. Complementation of strains was performed with strain harboring pSEVA224-*mxtR* or pSEVA224-*erdR* or the empty plasmid pSEVA224 and induced with 0.5 mM IPTG. Mean values and standard deviations of at least three biological replicates are displayed.

The growth of *P. alloputida* KT2440, Δ*mxtR* or Δ*erdR* in M9 minimal medium with succinate or glucose as a sole carbon source or complex King’s B (KB) medium was not affected by the deletion of *mxtR* or *erdR* (Figs. 2C and S1). Plasmid-based complementation of the two mutants restored growth on propionate, although delayed by a few hours for the *mxtR* complementation (Fig. 2D). The results indicate that *mxtR* and *erdR* are essential for utilizing propionate as a carbon source.

### 3.3. MxtR/ErdR affects the expression of genes encoding enzymes of the 2-MCC

The previously described target genes of MxtR/ErdR (Jacob et al., 2017; Sepulveda and Lupas, 2017) gave no indication that the two-component system is involved in the regulation of propionate metabolism. Concurrently, previous investigations have demonstrated that the utilization of propionate by *Pseudomonas* species requires activation to propionyl-CoA in conjunction with the 2-MCC (Ewering et al., 2006; Thompson et al., 2020; Cui et al., 2022). In *P. alloputida* KT2440, the propionyl-CoA synthetase is encoded by *prpE* (PP_2351), while the 2-MCC genes belong to the *prp* gene cluster, which comprises loci PP_2333 to PP_2339 (Fig. 3A). PP_2333 encodes a putative transcriptional regulator of the GntR protein family. Its *P. aeruginosa* ortholog was previously shown to act as a 2-methylisocitrate sensing repressor of the *prp* cluster and virulence associated genes (Cui et al., 2022). Genes PP_2333 and *mmgF* (PP_2334; 2-methylcitrate lyase) partially overlap (Fig. 3A) and are presumed to be co-transcribed from a promoter region upstream of PP_2333.

**Fig. 3:**
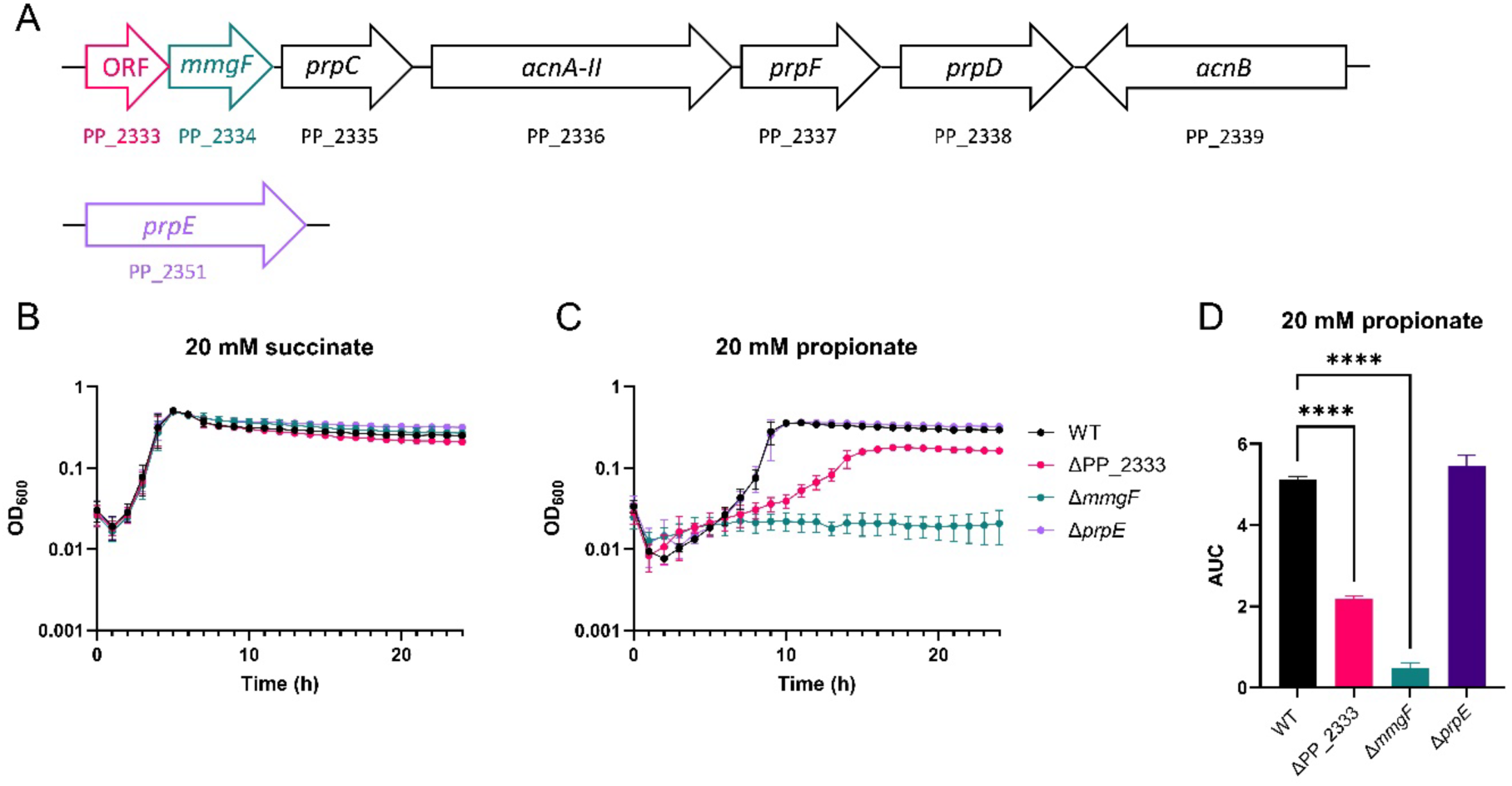
Importance of *prp* gene cluster in *P. alloputida* KT2440. (A) Schematic illustration of the *prp* gene cluster and *prpE* of *P. alloputida* KT2440. Scheme was created using genetic information provided by the Pseudomonas Genome Database (Winsor et al., 2016). (B, C) Growth analysis of wild type (WT), ΔPP_2333, Δ*mmgF* and Δ*prpE* in M9 minimal medium supplemented with 20 mM succinate or propionate. (D) Area under the curve (AUC) of WT, ΔPP_2333, Δ*mmgF* and Δ*prpE* growth curves in M9 minimal medium supplemented with 20 mM propionate. Growth analyses were performed with an initial OD_600_ of 0.1 and cells were cultivated in 96-well plates in a CLARIOstar Plus plate reader for 24 h, measuring OD_600_ hourly. Mean values and standard deviations of at least three biological replicates are displayed. \*\*\*\**p* < 0.0001 (ANOVA and Dunnett’s multiple comparison test).

We first assessed whether genes previously proposed to be involved in propionate metabolism are required for propionate utilization by *P. alloputida* KT2440 under our experimental conditions. To this end, we generated individual deletion mutants of PP_2333, *mmgF* (PP_2334), and *prpE* (PP_2351) and examined growth of the resulting strains on different carbon sources. None of the deletion mutants displayed a growth defect when succinate was provided as the carbon source (Fig. 3B). Deletion of *mmgF* abolished growth on propionate, demonstrating that this gene is essential for propionate utilization. In contrast, the PP_2333 deletion mutant retained the ability to grow on propionate but exhibited a reduced growth rate compared with the wild type. Deletion of *prpE*, however, had no significant effect on growth on propionate (Fig. 3C, D). The absence of a growth defect for strains on succinate suggested that the observed phenotypes were specific to propionate utilization. Expression of *mmgF* from the plasmid pUCP20-ANT2-MCS partially restored growth of the *mmgF* mutant on propionate, whereas plasmid-based expression of PP_2333 had no impact on the growth phenotype (Fig. S2). These results support an essential role of 2-MCC in propionate utilization and suggest that the absence of PrpE activity is compensated by other acyl-CoA synthetases, such as Acs (PP_3458), AcsA-I (PP_4487), and AcsA-II (PP_4702), as previously reported for other bacterial species (Palacios et al., 2003).

In a first attempt to obtain information on the mechanistic basis of the effect of MxtR and ErdR on propionate metabolism, we compared the expression of PP_2333, *mmgF* and *prpE* in the wild type with that of the *mxtR* and *erdR* mutants. For this purpose, the individual strains were pre-grown in M9 with succinate and transferred to either M9 with propionate or succinate. After one hour of incubation, the relative transcript levels of the three genes were determined by qRT-PCR. Transcript levels of each strain in propionate were compared to those in succinate.

In the wild-type strain, incubation with propionate resulted in an approximately 3.5-fold increase in the relative transcript levels of PP_2333 and *mmgF* compared with succinate. This induction was largely dependent on the presence of *mxtR* and *erdR* (Fig. 4A). In contrast, the relative transcript level of *prpE* in wild-type cells was approximately 10-fold lower following growth on propionate than on succinate. This reduction was independent of *mxtR* and *erdR* (Fig. 4A).

**Fig. 4:**
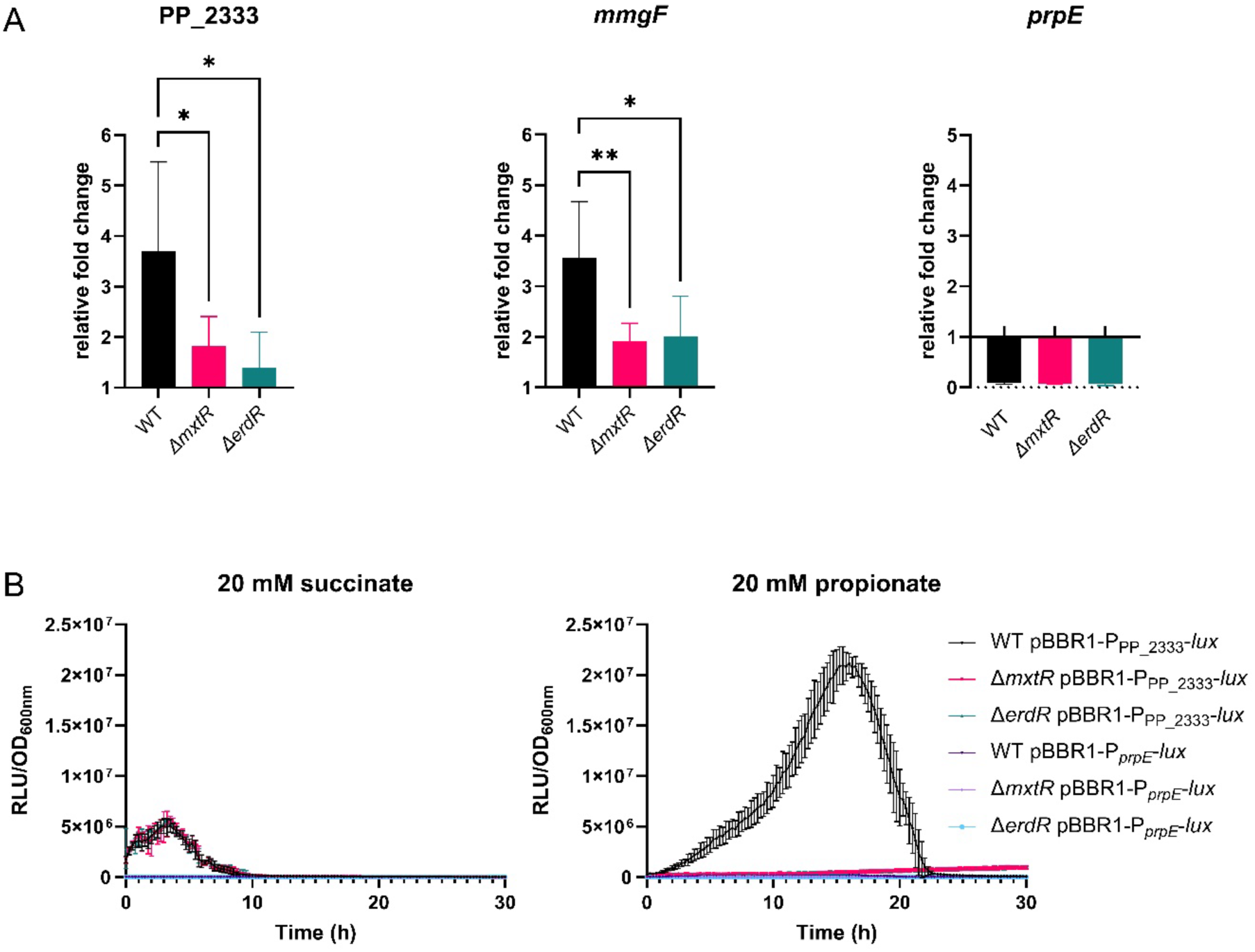
Expression levels of PP_2333, *mmgF* and *prpE* in wild-type, Δ*mxtR*, and Δ*erdR* cells. (A) Relative fold change of target genes of cells cultivated in M9 minimal medium with propionate compared to succinate determined via qRT-PCR. Cells were pre-cultured in M9 with 40 mM succinate, washed, shifted to M9 with either 40 mM succinate or propionate (start OD_600_ of ∼0.5) and cultivated for 1 h. Expression values were normalized to *gapA* (glyceraldehyde 3-phosphate dehydrogenase). (B) Promoter activity of PP_2333 and *prpE* promoters in wild-type (WT), Δ*mxtR*, and Δ*erdR* strains cultivated in M9 with 20 mM succinate or propionate, as measured in relative light units per OD_600_ (RLU/OD_600_). Growth and expression analyses were performed with an initial OD_600_ of 0.1 and cells were incubated in 96-well plates in a CLARIOstar Plus plate reader for 30 h, measuring the RLU and OD_600_ quarter-hourly. Mean values and standard deviations of at least three biological replicates are displayed. \**p* < 0.05 and \*\**p* < 0.01 (ANOVA and Dunnett’s multiple comparison test).

In addition to qRT-PCR, transcriptional expression was analyzed using reporter gene fusions. For this purpose, the nucleotide sequences upstream of PP_2333 and *prp* were fused to the *lux* gene cluster in pBBR1-MCS5-*lux* (Gödeke et al., 2011). The resulting plasmids were subsequently introduced into the wild-type, Δ*mxtR*, and Δ*erdR* strains. Since PP_2333 and *mmgF* are presumably co-transcribed from the same promoter region and partially overlap (Fig. 3A), no separate reporter fusion was constructed for *mmgF*. Reporter activity was assessed in M9 medium containing either succinate or propionate as the sole carbon source by measuring luminescence (RLU) and normalizing the values to the corresponding OD_600_.

During growth on succinate, low activity of the *P*_PP_2333_-*lux* transcriptional fusion was detected during the first 10 h and was independent of *mxtR* and *erdR*. In contrast, growth on propionate resulted in a significant increase in *P*_PP_2333_-*lux* activity in the wild-type strain over 20 h, and this induction was strictly dependent on *mxtR* and *erdR* (Figs. 4B and S3A). For the *P*_prpE_-*lux* fusion, wild-type cells grown on propionate exhibited a slight increase in reporter activity compared with cells grown on succinate. In contrast, no detectable luminescence was observed in the Δ*mxtR* or Δ*erdR* mutants under either growth condition (Figs. 4B and S3B).

Taken together, the qRT-PCR and transcriptional reporter fusion analyses indicate that, compared with succinate, growth on propionate stimulated expression of the PP_2333 and *mmgF* genes within the *prp* gene cluster. This induction was dependent on the presence of *mxtR* and *erdR*. In contrast, expression of *prpE* was only slightly affected by the cultivation conditions, and no detectable reporter activity was observed in the Δ*mxtR* and Δ*erdR* mutants.

### 3.4. ErdR binds a distinct DNA motif upstream of PP_2333 and *prpE*

Next, we tested the possibility whether the MxtR/ErdR TCS directly effects expression of genes of the 2-MCC gene cluster. For this purpose, the putative promoter regions upstream of PP_2333 and *prpE* were amplified by PCR, and ErdR was purified by Ni-NTA chromatography and gel filtration (Fig. S4). The resulting protein was phosphorylated with acetyl phosphate.

Binding of ErdR to the putative promoter regions was analyzed by EMSA. Addition of purified ErdR to the upstream regions of PP_2333 and *prpE* shifted both bands in the EMSA, indicating binding of ErdR to these regions. The shift was also observed for *scpC*, a previously identified direct target of ErdR (positive control), but was not detectable for an unrelated DNA region (*sodB*, negative control) (Fig. 5).

**Fig. 5:**
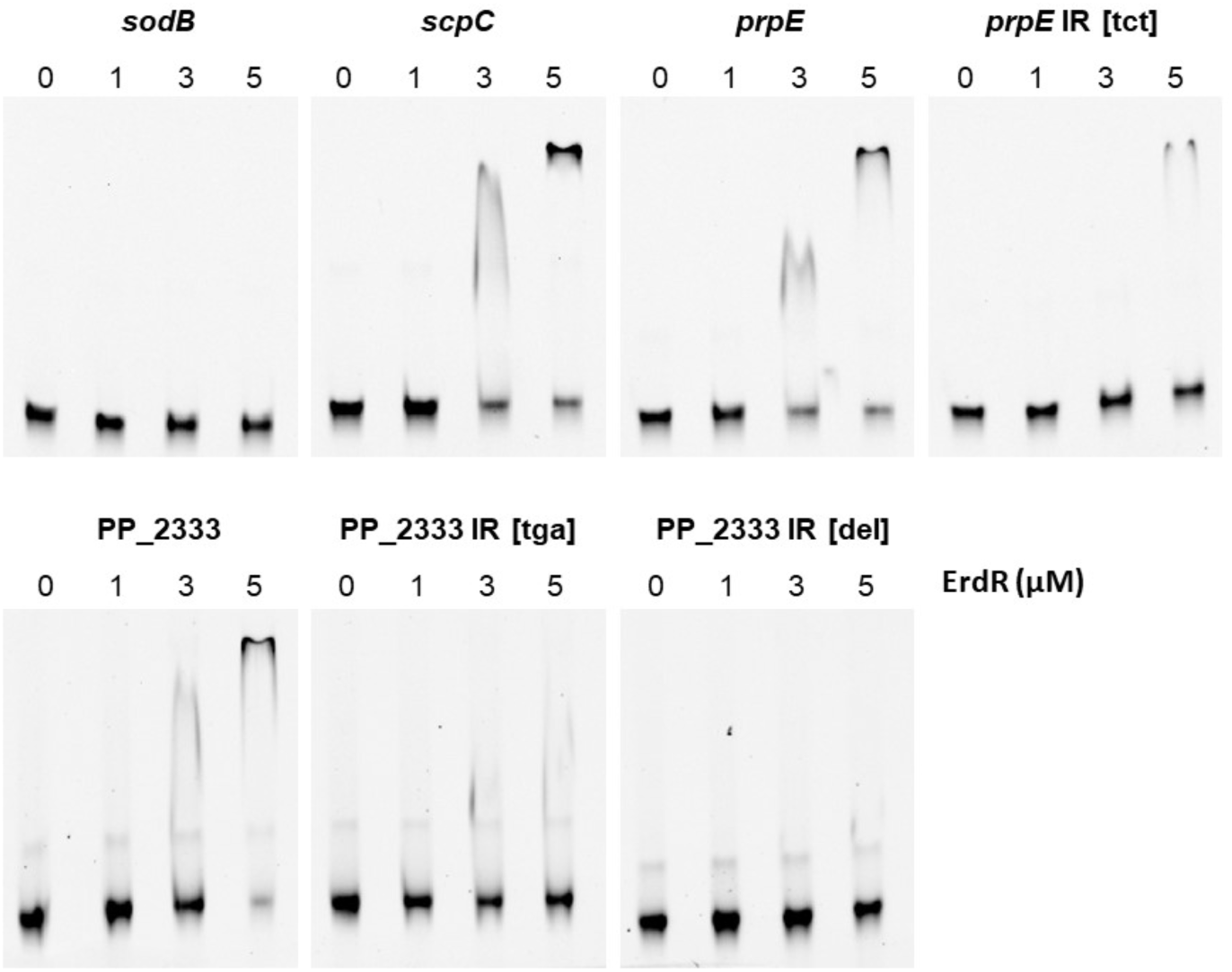
Binding of ErdR to the putative promoter regions of PP_2333 and *prpE*. In the EMSA, 1 nM of amplified and Cyanine5-tagged promoter regions (up to 160 bp upstream of respective start codon) was incubated with increasing concentrations of phosphorylated ErdR (0, 1, 3 or 5 μM). *P_scpC_* was used as a positive control, whereas an internal region of *sodB* acted as a negative control (Henriquez and Jung, 2021). A representative EMSA from three independent replicates is shown. *prpE* IR [tct], PP_2333 IR [tga], altered inverted repeat; PP_2333 IR [del], deleted inverted repeat.

In an effort to obtain more information on the sites of ErdR-binding regions and motifs in the upstream regions of PP_2333 and *prpE*, we analyzed the sequences upstream of PP_2333 and *prpE*. We identified the motif **GAC**NNNN**GTC** in the upstream regions of both PP_2333 and *prpE,* identical to the imperfect inverted repeat described previously for the ErdR homolog CrbR (**GAC**NNNN**GTC** or TACNNNNGTC) (Sepulveda and Lupas, 2017). To investigate whether ErdR does indeed bind to these sequences, the motif was altered in order to disrupt the imperfect palindrome. Thus, we mutated the PP_2333 motif from TTA**GAC**GA TT**GTC**GACA to TTA**GAC**GA TT**TGA**GACA and the *prpE* motif from TGT**GAC**TT TA**GTC**CCAT to TGT**GAC**TT TA**TCT**CCAT. In addition, the entire **GAC**NNNN**GTC** motifs were deleted.

EMSA analysis revealed that mutation of the inverted repeat in the *prpE* upstream region reduced ErdR binding, as indicated by the absence of a shifted band at 3 µM ErdR and a reduced band intensity at 5 µM ErdR. Similarly, disruption of the inverted repeat in the PP_2333 upstream region diminished ErdR binding. Notably, complete deletion of the inverted repeat abolished ErdR binding to the PP_2333 promoter region (Fig. 5).

In addition to EMSA, we investigated the effect of altering the putative promoter regions *P_prpE_* and *P*_PP_2333_ on gene expression using the *lux*-based reporter assay described above. In the wild-type strain grown on propionate, both disruption and deletion of the inverted repeat in *P*_PP_2333_ resulted in a significant decrease in *lux* expression compared with the promoter containing the intact inverted repeat (Fig. 6A, B). Alteration of the inverted repeat in the *prpE* upstream region also resulted in reduced expression (Fig. 6B). However, *prpE* expression levels were generally approximately 80-fold lower than those of PP_2333 (Fig. S5), which may limit the extent to which changes in expression can be reliably detected.

**Fig. 6:**
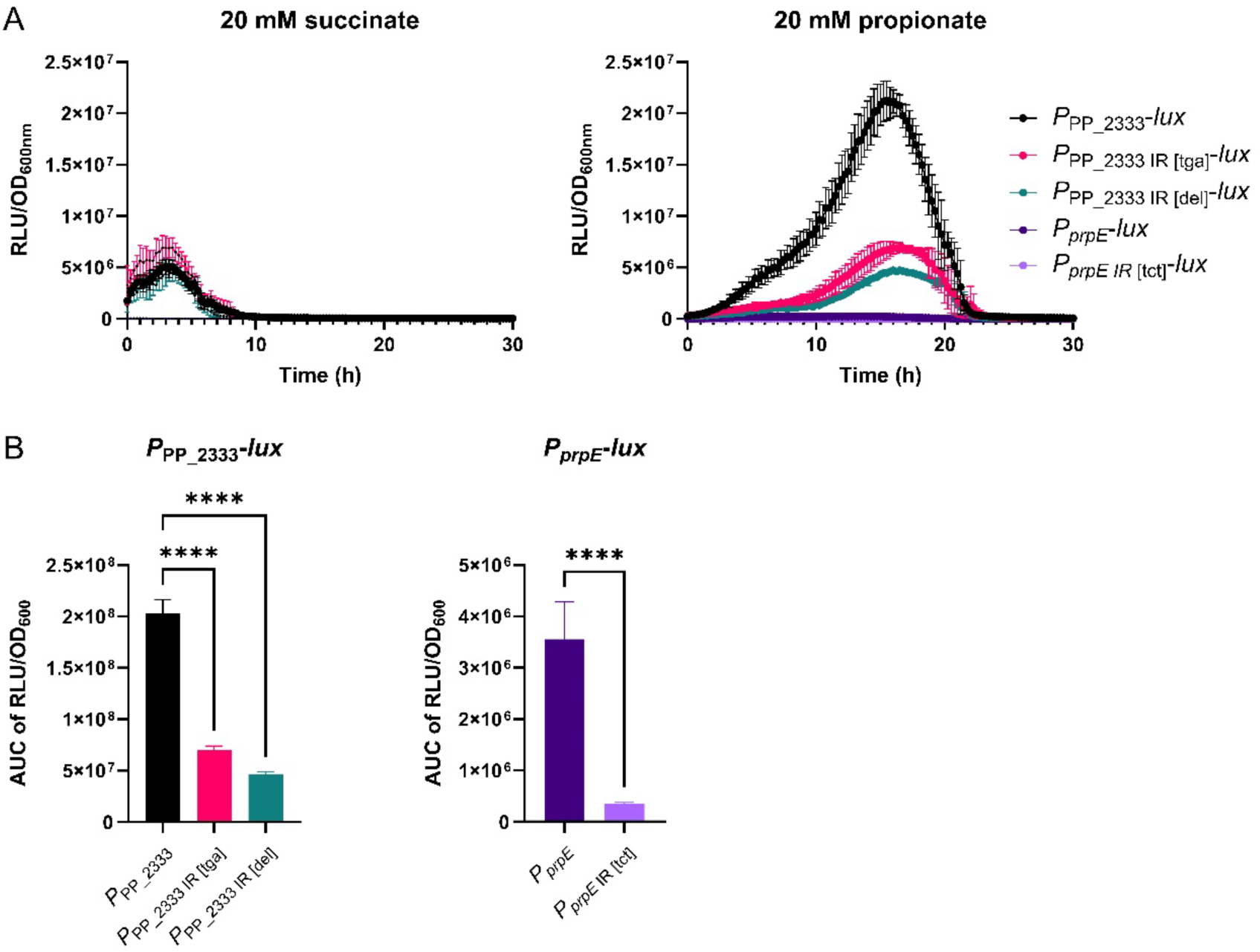
Impact of mutating putative promoter regions of PP_2333 and *prpE* on expression. (A) Unmodified and modified promoter activity of PP_2333 and *prpE* in the wild-type (WT) strain, cultivated in M9 with 20 mM succinate or propionate, represented as relative light units per OD600 (RLU/OD_600_). (B) AUC of RLU/OD_600_ of promoter variants of PP_2333 and *prpE* in the wild-type strain, cultivated in M9 with 20 mM propionate. Growth and expression analyses were performed with an initial OD_600_ of 0.1 and cells were cultivated in 96-well plates in a CLARIOstar Plus plate reader for 30 h, measuring OD_600_ and luminescence quarter-hourly. Mean values of at least three biological replicates are displayed. \*\*\*\**p* < 0.0001 (ANOVA and Dunnett’s multiple comparison test, t-test).

In the Δ*mxtR* and Δ*erdR* strains grown on propionate, mutation of the inverted repeats in *P*_PP_2333_ and *P_prpE_* did not result in significant changes in RLU/OD_600_ (Fig. S5). Furthermore, alteration of the promoter regions had no detectable effect on *lux* expression when cells were grown on succinate (Fig. 6A). Together with the EMSA results, these findings indicate that the identified inverted repeats contribute to ErdR-dependent regulation of PP_2333 and *prpE* expression under propionate-growth conditions.

### 3.5. Analysis of the impact of other direct or indirect target genes on growth on different carbon sources

To further assess the physiological relevance of the MxtR/ErdR regulatory network, we investigated further direct or indirect MxtR/ErdR target genes, as determined by RNAseq (Fig. 1A-B). Thus, we compared the growth of the wild type, Δ*mxtR* and Δ*erdR* strains with those of selected target gene deletion mutants on acetate, propionate, pyruvate and succinate as carbon sources.

Deletion of PP_0353, PP_0354, PP_2861, *yjcH*, or *actP-I* did not substantially affect growth on acetate or succinate (Fig. 7). On propionate, however, deletion of either PP_0353 or PP_0354 resulted in reduced growth, whereas deletion of PP_2861, *yjcH*, or *actP-I* had no significant effect. On pyruvate, growth was impaired in the ΔPP_0353, Δ*yjcH*, and Δ*actP-I* mutants, while deletion of the remaining genes caused little or no phenotype. In contrast, *mqo-I* proved essential for growth on acetate and propionate. Although the Δ*mqo-I* strain grew on succinate and pyruvate, the growth rate was considerably reduced compared to the wild-type strain (Fig. 7).

**Fig. 7:**
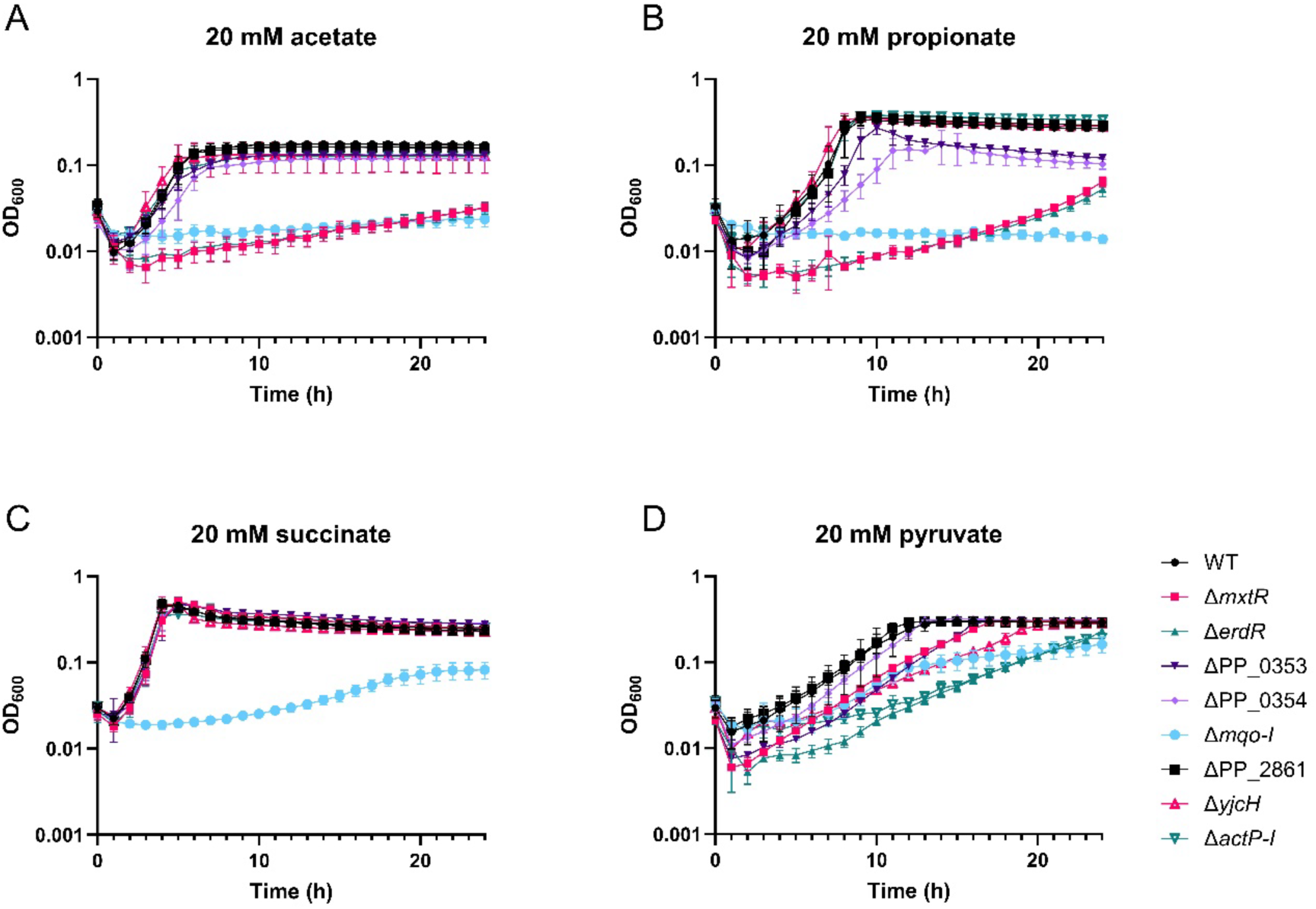
Phenotypic analyses of deletion mutants for suspected MxtR/ErdR target genes in minimal medium. Growth of wild-type (WT), Δ*mxtR*, Δ*erdR*, ΔPP_0353, ΔPP_0354, Δ*mqo-I*, ΔPP_2861, Δ*yjcH*, and Δ*actP-I* strains in M9 minimal medium supplemented with either 20 mM (A) acetate, (B) propionate, (C) succinate or (D) pyruvate as a sole carbon source. Growth analyses were performed with an initial OD_600_ of 0.1 and cells were cultivated in 96-well plates in a CLARIOstar Plus plate reader for 24 h, measuring OD_600_ hourly. Mean values and standard deviations of at least three biological replicates are displayed.

Next, we investigated whether purified ErdR binds to the putative promoter regions of *mqo-I*, *yjcH*, and PP_2861. EMSA analysis revealed a mobility shift for all three DNA fragments upon addition of ErdR, indicating that ErdR directly interacts with the corresponding upstream regions (Fig. 8) although a sequence with high similarity to the above-described imperfect inverted repeat was not identified upstream of *mqo-I* and PP_2861 (Fig. S6). The binding observed for *yjcH* is consistent with previous analyses of *actP-I* expression (Henriquez and Jung, 2021). In contrast, the mobility shifts observed for the *mqo-I* and PP_2861 upstream regions provide the first evidence that the MxtR/ErdR TCS may directly regulate genes associated with the TCA cycle and chemotaxis, respectively.

**Fig. 8:**
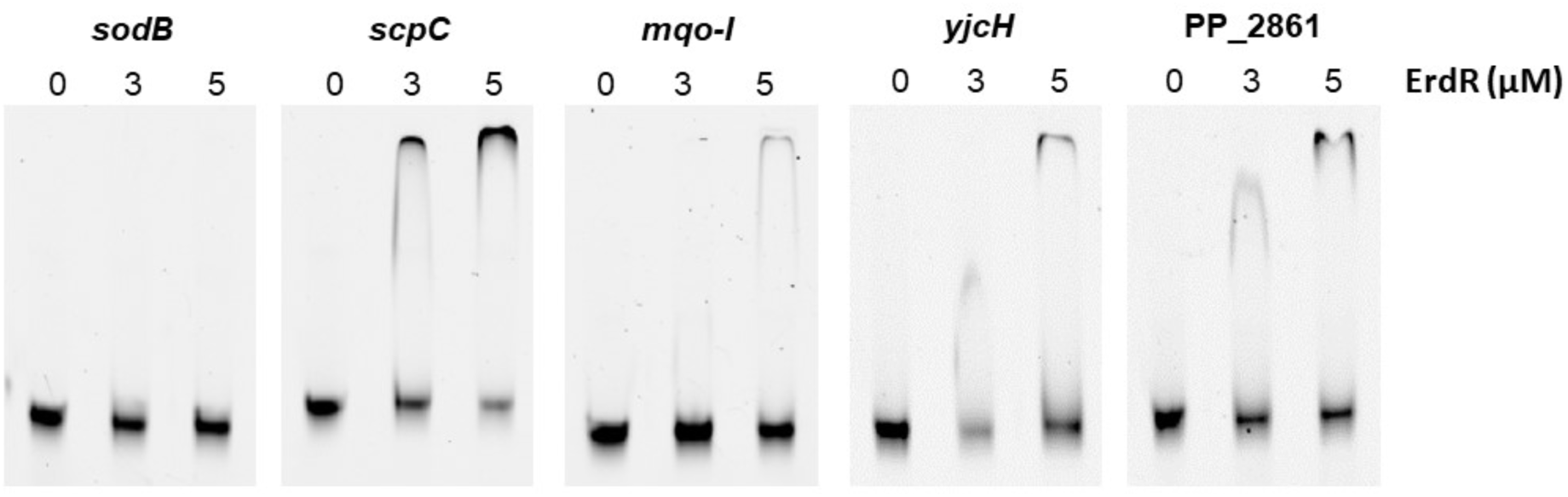
Binding of ErdR to upstream regions of *mqo-I*, *yjcH*, and PP_2861. In the EMSA, 1 nM of amplified and Cyanine5-tagged promoter regions (up to 160 bp upstream of respective start codon) was incubated with increasing concentrations of phosphorylated ErdR (0, 3, 5 μM). *P_scpC_* was used as a positive control, whereas an internal region of *sodB* acted as a negative control (Henriquez and Jung, 2021). A representative EMSA from three independent replicates is shown.

## 4. Discussion

The present study substantially expands the current understanding of the MxtR/ErdR two-component system in *P. alloputida* KT2440. Previous studies established MxtR/ErdR as the central regulator of acetate assimilation through activation of *acsA-I* and a limited number of genes involved in acetate transport and metabolism in *V. cholerae,* different *Pseudomonas species*, and *A. baumannii* (Jacob et al., 2017; Sepulveda and Lupas, 2017; Henriquez and Jung, 2021; Henriquez et al., 2023; Pokhrel et al., 2026). Our results demonstrate that the regulatory scope of MxtR/ErdR extends far beyond acetate utilization. Comparative transcriptomics, physiological analyses, promoter-reporter assays, mutational analyses and EMSAs collectively demonstrate an expanded regulon comprising genes involved in propionate utilization, central carbon metabolism, membrane transport, chemotaxis and signal transduction. Most importantly, our data identify propionate metabolism as a previously unrecognized physiological function of this conserved signaling system, indicating that MxtR/ErdR coordinates bacterial adaptation to multiple SCFA.

The transcriptomic analyses reveal that MxtR/ErdR primarily functions as a global transcriptional activator. In addition to confirming all previously identified targets (*acsA-I*, *actP-I*, *scpC* and PP_0354) (Henriquez and Jung, 2021; Henriquez et al., 2023), numerous additional genes involved in 2-MCC, TCA cycle, glyoxylate shunt, Ped pathway (oxidation of alcohols and aldehydes), solute transport and signal transduction were downregulated in the *mxtR-*H806N and ΔerdR strains. The broad metabolic response suggests that MxtR/ErdR functions upstream of several interconnected metabolic pathways rather than regulating acetate utilization in isolation. The hypothesis that MxtR/ErdR acts as a master regulator in a hierarchical system is supported also by previous analyses of the role of ErdR in the oxidation of alcohols (Görisch, 2003; Kretzschmar et al., 2010). Interestingly, the kinase mutant *mxtR-H806N* exhibited a much larger number of differentially expressed genes than the Δ*erdR* mutant. Since the changes in the additional genes detected in the *mxtR-*H806N strain are relatively minor compared to the most strongly differentially expressed genes and since they primarily affect general functions, such as ribosomal protein biosynthesis, oxidative phosphorylation, and flagellar biogenesis, these are likely indirect consequences of the altered metabolism. However, additional regulatory functions of MxtR for example in a sensor kinase network similar to the GacS-RetS network (Francis et al., 2018) cannot be excluded.

The most significant finding of this study is the discovery that MxtR/ErdR is essential for propionate utilization. Although acetate and propionate enter metabolism through distinct pathways, both substrates require activation to CoA esters before subsequent degradation. While this activation enables assimilation into central metabolism, excessive accumulation of acetyl-CoA or propionyl-CoA may disturb intracellular CoA homeostasis, inhibit CoA-dependent enzymes and alter global metabolic regulation through changes in acyl-CoA and acetyl-phosphate pools (Wolfe, 2005; Pinhal et al., 2019; Ren et al., 2019). It therefore appears physiologically advantageous that a single regulatory system coordinates transport, activation and subsequent degradation of both major SCFAs. Such regulation would permit rapid metabolic adaptation while preventing accumulation of potentially toxic intermediates.

A central finding of this work is the identification of the 2-MCC as a direct target of ErdR. The 2-MCC is widely conserved among Gram-negative bacteria and constitutes the principal pathway for detoxification and assimilation of propionyl-CoA generated during propionate degradation or β-oxidation of odd-chain fatty acids (Textor et al., 1997; Horswill and Escalante-Semerena, 1999; Claes et al., 2002). Besides serving as a carbon source, efficient degradation of propionyl-CoA is essential because this metabolite inhibits several CoA-dependent enzymes and interferes with central metabolism when allowed to accumulate (Horswill and Escalante-Semerena, 1999; Brock et al., 2002). Our physiological analyses demonstrating loss of growth on propionate in the Δ*mxtR* and Δ*erdR* mutants are therefore fully consistent with the observed failure to induce genes of the 2-MCC. Moreover, promoter-reporter analyses together with EMSAs reveal direct ErdR binding upstream of PP_2333 (a homolog of the *P. aeruginosa* gene *pmiR*, which encodes a transcriptional regulator sensing the 2-MCC intermediate 2-methylisocitrate (Cui et al., 2022) as the first gene of the 2-MCC gene cluster), demonstrating that regulation of the 2-MCC forms an integral component of the MxtR/ErdR regulon rather than representing an indirect metabolic consequence.

Interestingly, regulation of *prpE* appears more complex than regulation of the 2-MCC gene cluster. Although EMSAs demonstrate ErdR binding to the promoter region and disruption of the conserved inverted repeat reduces binding affinity, expression analyses failed to demonstrate a strong dependence on growth conditions and MxtR/ErdR. Furthermore, deletion of *prpE* did not impair growth on propionate. This observation is consistent with the remarkable redundancy of acyl-CoA synthetases encoded by *P. alloputida*, including several acetyl-CoA synthetases with overlapping substrate spectra (Nelson et al., 2002; Belda et al., 2016). Functional redundancy of propionyl-CoA activating enzymes has similarly been reported in *Salmonella enterica* and other bacteria (Horswill and Escalante-Semerena, 1999), suggesting that *prpE* contributes to optimization of propionate metabolism rather than determining its essentiality. Future metabolic flux analyses may clarify the quantitative contribution of the different acyl-CoA synthetases during growth on SCFAs.

The analysis of individual target genes further illustrates that the physiological consequences of MxtR/ErdR arise primarily through coordinated regulation and finetuning of multiple metabolic functions. Within this network, the function of individual direct MxtR/ErdR target genes remained unclear due to the absence of strong phenotypic defects in the corresponding deletion mutants. These include the genes PP_0353 and PP_0354, which are most strongly dependent on MxtR/ErdR and whose deletion resulted in only moderate inhibition of growth on propionate and pyruvate and no effect on acetate and succinate. The domain structure of PP_0354 (putative cyclic nucleotide-monophosphate (cNMP) binding domain, 2x CBS, nucleotidyltransferase) suggests an involvement in cNMP and energy (AMP, ATP) sensing as well as control of pathway activities by protein modification, similar to the mechanism described for the reversible lysine acetylation of acetyl-CoA synthetase (Zheng et al., 2025) or reversible uridylation of P_II_ (Atkinson et al., 1994). PP_0353 contains a putative exonuclease domain similar to the one found in RNase-T, which can affect tRNA and rRNA maturation (Zuo and Deutscher, 2002), but also cleave DNA. Further protein chemical and metabolic analyses are necessary to understand the role of the relevant proteins for the metabolism of SCFA. Regardless, this observation highlights the importance of regulatory integration rather than individual gene essentiality. The impaired growth of Δ*actP-I* and Δ*yjcH* on pyruvate confirms previous work demonstrating that MxtR/ErdR contributes to pyruvate utilization through regulation of substrate uptake (Henriquez et al., 2023).

In contrast, deletion of *mqo-I* abolished growth on acetate and propionate and impaired growth on succinate and pyruvate, highlighting the central role of malate oxidoreductase in respiratory metabolism and energy conservation (Kretzschmar et al., 2002; Mellgren et al., 2009). The demonstrated binding of ErdR to the putative *mqo-I* promoter region provides initial evidence that MxtR/ErdR may directly influence carbon flux through the TCA cycle by regulating gene expression. However, although *mqo-I* expression is modulated by MxtR/ErdR, the physiological function of the encoded enzyme extends beyond this regulatory network.

Another intriguing observation is the coordinated regulation of metabolic genes together with genes involved in environmental sensing. Among these, the recently characterized chemoreceptor PP_2861, which mediates taxis towards acetate and propionate (Garcia et al., 2015), represents a particularly interesting candidate. Although deletion of PP_2861 did not influence growth in well-mixed liquid culture, simultaneous regulation of chemotaxis and catabolism may allow *P. alloputida* to couple movement towards favorable carbon sources in a more structured environment. Such integration between chemotactic behavior and metabolism is increasingly recognized as an important determinant of ecological fitness in environmental pseudomonads inhabiting nutrient gradients within soil and the rhizosphere (Matilla and Krell, 2018).

Our promoter analyses further support the emerging model of ErdR-dependent transcriptional regulation. Mutation or deletion of the conserved imperfect inverted repeat significantly reduced promoter activity and impaired ErdR binding, confirming previous predictions that this sequence constitutes the ErdR recognition motif (Sepulveda and Lupas, 2017). The presence of this motif in both previously characterized and newly identified target promoters suggests that it represents a conserved regulatory signature across the MxtR/ErdR regulon. Nevertheless, the differing promoter responses observed for PP_2333 and *prpE* indicate that promoter architecture, accessory transcription factors or metabolic effectors probably modulate ErdR-dependent activation.

Finally, the nature of the signal perceived by MxtR remains one of the most intriguing unresolved questions. Unlike canonical sensor kinases, MxtR contains an N-terminal sodium:solute symporter (SSS) domain. This unusual architecture resembles that of CbrA, another SSS-containing sensor kinase that coordinates carbon and nitrogen metabolism in pseudomonads (Nishijyo et al., 2001; Valentini et al., 2014; Wirtz et al., 2020). Screening using a thermoshift assay for potential ligands of the sensor kinase AcmR (an orthologue of MxtR in *A. baumannii*) showed that various SCFAs, such as acetate, propionate, butyrate and valerate, affect the protein’s thermostability. The authors hypothesized that acetate is the physiological signal for AcmS (Pokhrel et al., 2026). Given that MxtR regulates both acetate and propionate metabolism, it appears unlikely that the kinase responds exclusively to one individual substrate. Instead, MxtR with its SSS and PAS domains may detect a shared physiological consequence of SCFA assimilation, such as changes in intracellular acyl-CoA pools, CoA availability, membrane energetics or proton motive force. Elucidating the molecular mechanism of signal perception by the SSS domain therefore represents an important objective for future studies.

Collectively, our results establish MxtR/ErdR as considerably more than an acetate-switch regulator and we thus present the following model, proposing the MxtR/ErdR TCS as a master regulator of SCFA metabolism in *Pseudomonas* species and other Gammaproteobacteria (Fig. 9).

**Fig. 9:**
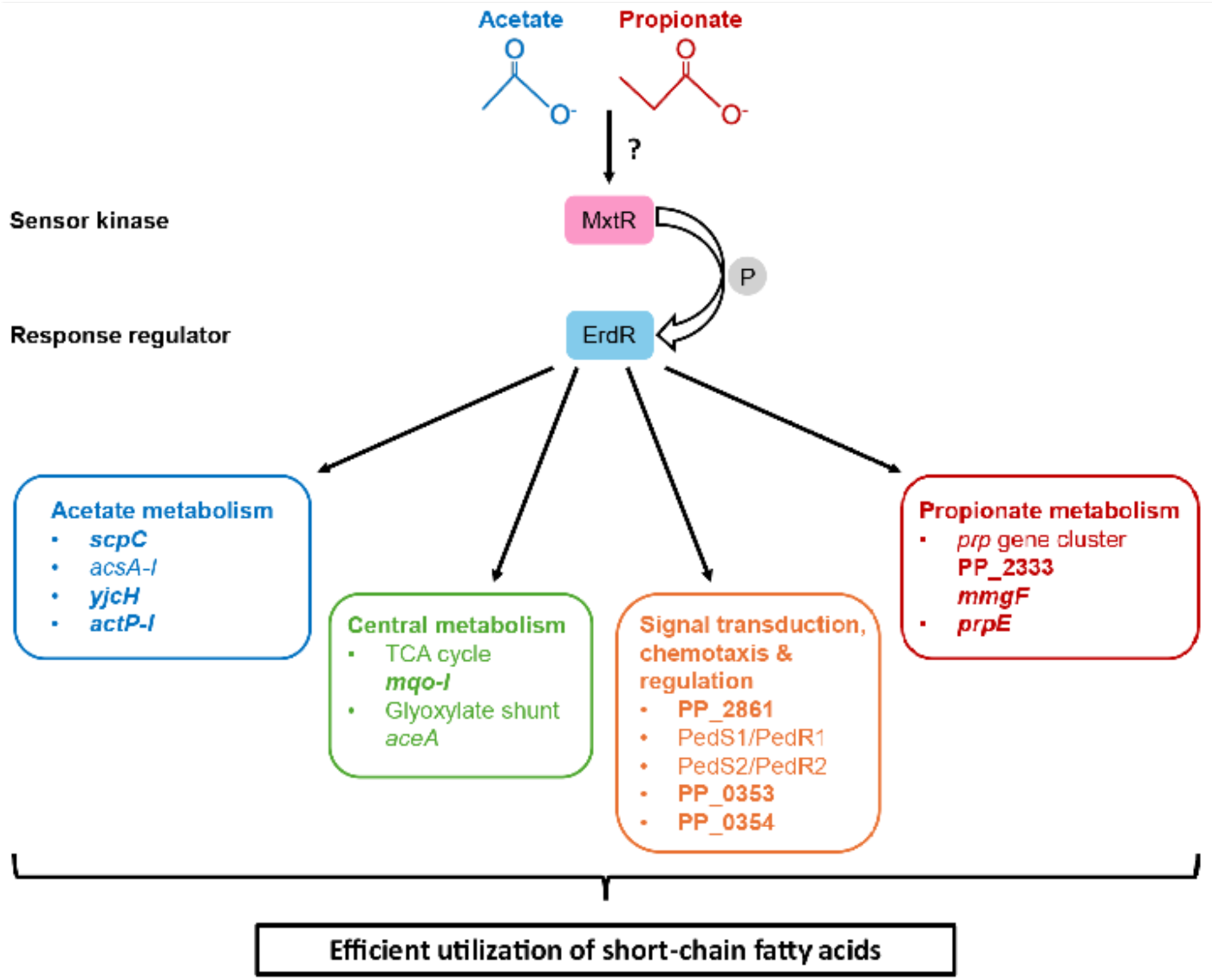
Model of the MxtR/ErdR regulatory network controlling SCFA metabolism in *P. alloputida* KT2440. MxtR is proposed to sense signals associated with the availability and/or metabolism of SCFAs, such as acetate and propionate. First evidence for acetate as the physiological ligand of the sensor kinase comes from the MxtR ortholog AcmS of *A. baumannii* (Pokhrel et al., 2026). MxtR then activates the response regulator ErdR through phosphorylation. The precise mechanism by which phosphotransfer occurs from the REC domain-containing MxtR to the REC domain of ErdR remains unknown. The activated ErdR protein directly induces the expression of genes necessary for the utilization of acetate and propionate (Henriquez and Jung, 2021; Henriquez et al., 2023), this study). EMSA analyses also suggest the direct regulation of selected genes involved in central carbon metabolism, regulation, and chemotaxis (this study). This coordinated regulation promotes the efficient utilization of acetate, propionate, and pyruvate, thereby maintaining SCFA homeostasis. **Bold** genes, binding of ErdR to promoter regions verified by EMSAs; other genes, directly or indirectly regulated by MxtR/ErdR.

## 5. Conclusion

In summary, we believe MxtR/ErdR to be a master regulator of SCFA metabolization, extending vastly beyond its role as a simple acetate switch, but rather being involved in the coordination of multiple metabolites at once, including propionate, acetate and presumably also a yet unknown stimulatory signal. The signaling system, with MxtR at the top of the hierarchy, coordinates a broader adaptive program that integrates signal transduction, transport, central carbon metabolism, chemotaxis and the metabolism of multiple SCFAs. Direct regulation of the 2-MCC provides a mechanistic link between acetate and propionate assimilation and suggests that MxtR/ErdR functions as a central regulator of SCFA homeostasis in *P. alloputida*. Because homologous signaling systems are widespread among Gammaproteobacteria, these findings likely have implications beyond pseudomonads and provide a framework for understanding how metabolically versatile bacteria coordinate carbon source availability with physiological adaptation.

## Acknowledgments

We thank Vera Wanat for her assistance in the construction of the deletion strains ΔPP_2861, Δ*mqo-I* and Δ*yjcH*. Furthermore, we thank Daniel Rehm for the construction of the Δ*mmgF* strain. Research in the group of H.J. was supported by the Deutsche Forschungsgemeinschaft, project JU333/6-2, and the Faculty of Biology, LMU Munich. In addition, this work was supported by the Luxembourg National Research Fund (FNR) (Project Code: 18865049).

## Supporting Material

Additional material can be found in the online version of this article at the publisher’s web-site:

## Supporting material

Table S1: Strains generated and used in this investigation.

Table S2. Plasmids generated and/or used during this investigation.

Table S3: Oligonucleotides used in this study.

Table S4: Genes downregulated in *mxtR*-H806N compared to the WT.

Table S5: Genes upregulated in *mxtR*-H806N compared to the WT.

Table S6: Genes downregulated in Δ*erdR* compared to the WT.

Table S7: Genes upregulated in Δ*erdR* compared to the WT.

Fig. S1: Impact of *mxtR* and *erdR* on growth in minimal or complex medium.

Fig. S2: Complementation of ΔPP_2333 and *prpE* of the *prp* gene cluster.

Fig. S3: Comparison of PP_2333 and *mmgF* expression levels in different carbon sources.

Fig. S4: Purification of 10H-ErdR via affinity and size exclusion chromatography (SEC).

Fig. S5: Impact of mutation of upstream promoter regions of PP_2333 and *prpE* on expression.

Fig. S6: Putative DNA binding sites of ErdR of *P. alloputida* KT2440.

