## Supplementary Material for "MxtR/ErdR is a central regulator of short-chain fatty acid metabolism in *Pseudomonas alloputida*"

[1]Present address: Università degli Studi di Siena, Siena, Italy

**Table S1: Strains generated and used in this investigation.**

| Strain | Description | Origin |
|---|---|---|
| *Pseudomonas alloputida* KT2440 | wild-type strain | (Bagdasarian et al., 1981; Nelson et al., 2002) |
| *Pseudomonas alloputida* KT2440 Δ*mxtR* | Δ*mxtR* (PP_1695) | (Henriquez and Jung, 2021) |
| *Pseudomonas alloputida* KT2440 Δ*erdR* | Δ*erdR* (PP_1635) | (Henriquez and Jung, 2021) |
| *Pseudomonas alloputida* KT2440 *mxtR*-H806N | Insertion of a point mutation in *mxtR* causing a histidine to asparagine substitution at residue 806 of MxtR | This work |
| *Pseudomonas alloputida* KT2440 ΔPP_2333 | ΔPP_2333 | This work |
| *Pseudomonas alloputida* KT2440 Δ*mmgF* | Δ*mmgF* (ΔPP_2334) | This work |
| *Pseudomonas alloputida* KT2440 Δ*prpE* | Δ*prpE* (ΔPP_2351) | This work |
| *Pseudomonas alloputida* KT2440 ΔPP_0353 | ΔPP_0353 | (Henriquez et al., 2023) |
| *Pseudomonas alloputida* KT2440 ΔPP_0354 | ΔPP_0354 | (Henriquez et al., 2023) |
| *Pseudomonas alloputida* KT2440 ΔPP_2861 | ΔPP_2861 | This work |
| *Pseudomonas alloputida* KT2440 Δ*mqo-I* | Δ*mqo-I* (ΔPP_0751) | This work |
| *Pseudomonas alloputida* KT2440 Δ*yjcH* | Δ*yjcH* (ΔPP_1742) | This work |
| *Pseudomonas alloputida* KT2440 Δ*actP-I* | Δ*actP-I* (ΔPP_1743) | (Henriquez et al., 2023) |
| *Escherichia coli* C41 | Also known as C41(DE3), derivative of BL21(DE3), used for protein overexpression | (Miroux and Walker, 1996) |
| *Escherichia coli* DH5αλpir | Cloning strain, *pir*$^+$, for propagation of R6K ori plasmids | (Miller and Mekalanos, 1988) |
| *Escherichia coli* WM3064 | DAP-auxotrophic conjugation donor strain, *pir*$^+$, RP4 transfer functions | W. Metcalf, unpublished data |

**Table S2: Plasmids generated and/or used during this investigation.**

| Plasmid | Description | Origin |
|---|---|---|
| pNPTS138-R6KT | Suicide plasmid for in-frame deletions; *mobRP4*$^{+}$,*oriR6K sacB* Km$^{r}$ | (Lassak et al., 2010) |
| pSEVA224 | Derivative of pSEVA221 with *lacI*$^{q}$*/Ptrc* expression system, Km$^{r}$ | (Silva-Rocha et al., 2013) |
| pSEVA224-*mxtR6H* | pSEVA224 derivative expressing C-terminally $His_6$-tagged MxtR (PP_1695) | This work |
| pSEVA224-10HFXGS-*erdR* | pSEVA224 derivative expressing N-terminally $His_{10}$-FXa-GS-tagged ErdR (PP_1635) | This work |
| pUCP20-ANT2-MCS | Anthranilate-inducible expression vector with multiple cloning site, Km$^{r}$ | (Hoffmann et al., 2021) |
| pUCP20-ANT2-PP_2333 | pUCP20-ANT2 derivative expressing PP_2333 | This work |
| pUCP20-ANT2-*mmgF* | pUCP20-ANT2 derivative expressing *mmgF* (PP_2334) | This work |
| pBBR1-MCS5-*luxCDABE* | pBBR1-MCS5 derivative containing promoterless *luxCDABE* | (Gödeke et al., 2011) |
| pBBR1-$P_{PP\_2333}$-*lux* | Derived from pBBR1-MCS5-*lux*, insertion of promoter region upstream of PP_2333 | This work |
| pBBR1- $P_{PP\_2333-IR-[tga]}$-*lux* | Mutation of 3 bases (GTC→TGA) in the upstream region of the start codon of PP_2333 | This work |
| pBBR1- $P_{PP\_2333-IR-[del]}$-*lux* | Deletion of imperfect inverted repeat in the upstream region of the start codon of PP_2333 | This work |
| pBBR1-$P_{prpE}$-*lux* | Derived from pBBR1-MCS5-*lux*, insertion of promoter region upstream of *prpE* (PP_2351) | This work |
| pBBR1- $P_{prpE-IR-[tct]}$-*lux* | Mutation of 3 bases (GTC→TCT) in the upstream region of the start codon of *prpE* (PP_2351) | This work |
| pET16b-*erdR* | Derivative of pET16b containing *erdR* (PP_1635) | (Henriquez and Jung, 2021) |

**Table S3: Oligonucleotides used in this investigation.**

| Primer designation | Sequence [5' to 3'] | Purpose |
|---|---|---|
| 1695_H806N_as | CATCAGGTCATTGCTGAC GGCGGC | Genomic *mxtR*-H806N substitution |
| 1695_H806N_s | GCCGTCAGCAATGACCT GATGCAA | |
| SubH806N_A_s_BamHI | TGGCCATGAGGATCCTTC CAGCGGATG | |
| SubH806N_B_as_NheI | CCATCGAAGGCTAGCACA TGCAACTGG | |
| Del1695_check_s | AAGGGCCGCGAATACCA GAC | |
| Del1695_check_as | ATCGTCGGCTGGTGTGCT TTAC | |
| delPP2333_As_SpeI | AGGTAGAAGacTAgtCA GCTTTCGCCATTG | Genomic deletion of PP_2333 |
| delPP2333_Aas_OL_new | CATTTCTCACCTCGTCTG CATATTGTCGACACC | |
| delPP2333_Bs_OL_new | GGTGTCGACAATATGCAG ACGAGGTGAGAAATG | |
| delPP2333_Bas_EcoRI | TTCGGTGATGaaTTCCGG GAAGATCATG | |
| delPP2333_check_s | AAGGCTTGGGTCAACCA GTG | |
| delPP2333_check_as | TACACGTTCTCGGCTGCT TTGTTC | |
| delPP2334_fragA_as OL | GAAATGACAGTCAAGCG CATGGAGCTTTAC | Genomic deletion of *mmgF* (PP_2334) |
| delPP2334_fragA_s_PstI | ATATCGCCCCCtGCAGCA AGATTTCCG | |
| delPP2334_fragB_as_Bam HI | GAAGACACCTTGGGatcC CACTTCCTG | |
| delPP2334_fragB_s_OL | GAAATGACAGTCAAGCG CATGGAGCTTTAC | |
| delPP2334_check_as | GACTTGTACTCCTGCGTT ACCG | |
| delPP2334_check_s | CGATCTGACAGCGTTGAA GTGG | |
| delPP2351_As_SpeI | GCTTCTCTTCCATCAACTa gTCCACCTGCAG | Genomic deletion of *prpE* (PP_2351) |
| delPP2351_Aas_OL | GTCAAACCATCAACCCGC GTAGCTCATGTCGTTAG | |
| delPP2351_Bs_OL | CTAACGACATGAGCTACG CGGGTTGATGGTTTGAC | |
| delPP2351_Bas_EcoRI | CTGTATCACGAACGaaTT CTGCCGGT | |
| delPP2351_check_s | TTGGTCGAGTATGAACTG CAG | |
| delPP2351_check_as | TGCTGTGCTTCGACGAAT TC | |
| mxtR_BamHI_s | ATCCAGGAGATggAtccAT GtCGTTGTCC | Complementation plasmid pSEVA224-*mxtR6H* |

| | | |
|---|---|---|
| seq_pSEVA224_as | GACTAGTCGCCAGGGTTTTCCCAGTCACG | |
| pUCP20-ANT2-10HFXGS_EcoRI_s | CTAGTAGGAGATGAATTCATGCATCATC | Complementation plasmid pSEVA224-10HFXGS-*erdR* |
| pUC19s | AAGTTGGGTAACGCCAGGGT | |
| pp_2333_NdeI_fw | TGTCGACCATATGCAGGACCTTTCCACC | Complementation plasmid pUCP20-ANT2-PP_2333 |
| pp_2333_PstI_rew | GGGTGCTCTCTGCAGTCATTTCTCACC | |
| pp_2334_NdeI_fw | ACGAGGTGACATATGACA | Complementation plasmid pUCP20-ANT2-*mmgF* |
| pp_2334_XbaI_rew | GGCGCTCTAGATTAGCCCTTCTTCTG | |
| Ppp2333_XhoI_rv | GGGCTCGAGAGGTCCTGCATATTG | PP_2333 promoter-*luxCDABE* reporter construction |
| Ppp2333_BamHI_fw | TTGACGCCAGGATCCCG | |
| InvRep_Ppp2333-tga_s | GTTCGCGATTAGACGATTTGAGACAATGTTC | |
| InvRep_Ppp2333-tga_as | GAACATTGTCTCAAATCGTCTAATCGCGAAC | |
| InvRep_PP2333-del-fw | ATGTTCATTTTGTCTTTGA | |
| InvRep_PP2333-del-rv | CGCGAACTATCACCAC | |
| Ppp2351 (prpE)_BamHI_fw | TCTAGAGGCGGATCCCCGTTATGGG | *prpE* promoter-*luxCDABE* reporter construction |
| Ppp2351 (prpE)_XhoI_rv | GTAGCTGTGCTCGAGGCTCATGTCGTTAG | |
| InvRep_Ppp2351-tct_s | GGTCGGTTGTGACTTTAT | |
| InvRep_Ppp2351-tct_as | GCCATGGAGATAAAGTCA | |
| gapA fw v2 qPCR | TACAACGCTCTGCCATTG | qRT-PCR |
| gapA rv v2 qPCR | TTGTCCAGCATCCGGT | |
| pp_2333 fw v1 qPCR | AGTTCTCTGCCACGCCTA | |
| pp_2333 rv v1 qPCR | CTCGTGTGCTGTTGTTGT | |
| pp_2334 fw v2 qpCR | CGCTGTACACAACCGAAG | |
| pp_2334 rv v2 qPCR | ACATTCTTTTGCGTGCC | |
| pp_2351 fw v2 qPCR | GTAAGCCCAAGGGTATC | |
| pp_2351 rv v2 qPCR | GCTTGCCTTCGTAGAACA | |
| PscpCF | GATACGATCACGGTACAT | Promoter amplification for EMSA |
| ScpcF EMSA | [Cyanine5]-TCTGCCGGGGCTTTTCTTT | |
| sodBF EMSA | [Cyanine5]-AACACCTATGTCGTGAACCT | |
| sodB R | TTCCAGTAGAAGGTGTGGTT | |
| pp_2333 F EMSA | [Cyanine5]-CGCGATCTGACAGCGTT | |
| pp_2333 R | AGGTCCTGCATATTGTCGA | |

| | | |
|---|---|---|
| pp_2351 EMSA F | [Cyanine5]-<br>CCGTTATGGGGCGTAATT | |
| pp_2351 R | GCTCATGTCGTTAGAAAC | |
| delPP0751_A_s_PstI | AAACCCTGTACCTGCAGT<br>TTCTGACTTAC | Genomic deletion of *mqo-I* (PP_0751) |
| delPP0751_A_as_OL | CGTTAAATGGCGCAGTAC<br>GCTTAAGCCGGTC | |
| delPP0751_B_s_OL | GACCGGCTTAAGCGTACT<br>GCGCCATTTAACG | |
| delPP0751_B_as_EcoRI | CCGATTGACGAATTCAAC<br>TTCACCGCTTC | |
| delPP0751_check_s | TGAAAGGCAAGCCGCTG<br>AGC | |
| delPP0751_check_as | CATCGACTGGGTGCAGAT<br>CAGC | |
| delPP1742_A_s_PstI | GACGCAGGCTGCAGTGT<br>GATGCTCAAG | Genomic deletion of *yjcH* (PP_1742) |
| delPP1742_A_as_OL | GGCGAATCATTGTTGGTC<br>GTTCATTGTTTTTG | |
| delPP1742_B_s_OL | CAAAAACAATGAACGACC<br>AACAATGATTCGCC | |
| delPP1742_B_as_EcoRI | GAACAGCACGTGAATTCA<br>CATCAGGATACC | |
| delPP1742_check_s | TGGGCCAAGAACAAGGA<br>CAACG | |
| delPP1742_check_as | CCTTGATCGCTTCGGAGA<br>ACAGC | |
| delPP2861_A_s | ATAGACGATGTCGACCAC<br>GATCCACAG | Genomic deletion of PP_2861 |
| delPP2861_A_as_OL | CTGATGGATGAACACTCG<br>GGTCTGAGCCTAC | |
| delPP2861_B_s_OL | GTAGGCTCAGACCCGAG<br>TGTTCATCCATCAG | |
| delPP2861_B_as_BamHI | ACGGTGACGGGATCCAA<br>CAGGTATTCG | |
| delPP2861_check_s | ACTTTCATTGCTTGCGGC<br>GCACTC | |
| delPP2861_check_as | TTTTCAGCGCATGGCTCC<br>AG | |

**Table S4: Genes downregulated in *mxtR*-H806N compared to the wild type.** Differentially expressed genes with $log_2$foldchange <-1 and adjusted *p*-value (padj) <0.05. Gene annotations/functions were retrieved from the *Pseudomonas* Genome Database (Winsor et al., 2016) and UniProt (UniProt, 2026).

| | **Gene / locus tag** | **$log_2$FoldChange** | **padj** | **Annotated function** |
|---|---|---|---|---|
| 1 | PP_0354 | -6.55 | 2.76E-229 | CBS domain-containing protein |
| 2 | PP_0353 | -6.29 | 3.03E-173 | Exonuclease |
| 3 | *yjcH* | -6.11 | 5.35E-60 | Inner membrane protein |
| 4 | PP_5559 | -4.19 | 2.41E-35 | Uncharacterized protein |
| 5 | *acsA_I* | -4.11 | 3.39E-84 | Acetyl-CoA synthetase |
| 6 | *pqqD_II* | -4.06 | 3.30E-66 | Coenzyme PQQ synthesis protein D |
| 7 | PP_2861 | -3.91 | 1.13E-83 | Methyl-accepting chemotaxis protein McpP |
| 8 | *actP_I* | -3.81 | 7.55E-79 | Acetate permease, Cation/acetate symporter ActP |
| 9 | PP_1587 | -3.62 | 3.96E-37 | NhaP-type Na+(K+)/H+ antiporter |
| 10 | *sthA* | -3.53 | 6.21E-24 | Soluble pyridine nucleotide transhydrogenase |
| 11 | PP_2663 | -3.48 | 6.58E-33 | GfdT protein |
| 12 | *scpC* | -3.32 | 2.55E-21 | Acetate:succinate CoA transferase |
| 13 | PP_1635 | -3.31 | 2.84E-48 | Response regulator ErdR |
| 14 | PP_3179 | -3.30 | 1.25E-48 | LysR family transcriptional regulator |
| 15 | PP_5297 | -3.29 | 1.95E-46 | Amino acid transporter |
| 16 | *mqo_I* | -3.28 | 1.20E-50 | Malate:quinone oxidoreductase |
| 17 | *exaE* | -2.76 | 2.46E-28 | PedR2 |
| 18 | PP_2664 | -2.70 | 1.15E-32 | Sensor histidine kinase |
| 19 | PP_0333 | -2.62 | 1.30E-18 | Retropepsin-like aspartic endopeptidase RimB domain-containing protein |
| 20 | PP_0051 | -2.57 | 8.06E-16 | Sigma-54 dependent transcriptional regulator |
| 21 | PP_5410 | -2.55 | 3.36E-02 | DeoR family transcriptional regulator |
| 22 | PP_4975 | -2.48 | 1.25E-48 | Long-chain acyl-CoA thioester hydrolase family protein |
| 23 | PP_3014 | -2.45 | 1.86E-26 | Lipoprotein |
| 24 | PP_2671 | -2.39 | 2.64E-25 | Histidine kinase PedS2 |
| 25 | *cti* | -2.39 | 1.40E-22 | Esterified fatty acid cis/trans isomerase |
| 26 | *opdN* | -2.38 | 5.07E-17 | Outer-membrane porin D |
| 27 | *pctA* | -2.27 | 2.45E-10 | Methyl-accepting chemotaxis protein PctA |
| 28 | PP_4677 | -2.22 | 4.52E-07 | CDP-diacylglycerol--serine O-phosphatidyltransferase |
| 29 | PP_4732 | -2.11 | 4.57E-17 | Coenzyme Q-binding protein COQ10 START domain-containing protein |
| 30 | PP_1832 | -2.10 | 1.31E-07 | Oxidase |

| | | | | |
|---|---|---|---|---|
| 31 | *benR* | -2.10 | 8.85E-13 | BenABC operon transcriptional activator |
| 32 | PP_2144 | -2.08 | 6.62E-09 | TetR family transcriptional regulator |
| 33 | PP_2667 | -2.06 | 1.64E-04 | PedC |
| 34 | *ccoN_II* | -2.05 | 2.27E-10 | Cbb3-type cytochrome c oxidase subunit 1 |
| 35 | PP_5620 | -2.04 | 9.08E-14 | MAE-28990/MAE-18760-like HEPN domain-containing protein |
| 36 | *dgkA_I* | -1.88 | 6.24E-14 | Diacylglycerol kinase |
| 37 | *aceA* | -1.85 | 3.14E-22 | Isocitrate lyase |
| 38 | PP_5659 | -1.84 | 1.76E-19 | Uncharacterized protein |
| 39 | PP_2446 | -1.83 | 1.56E-13 | Uncharacterized protein |
| 40 | *rnt* | -1.82 | 1.67E-02 | Ribonuclease T |
| 41 | PP_4254 | -1.75 | 3.16E-07 | Uncharacterized protein |
| 42 | PP_1157 | -1.68 | 1.11E-02 | Acetolactate synthase |
| 43 | *betI* | -1.65 | 6.32E-09 | HTH-type transcriptional regulator BetI |
| 44 | PP_5538 | -1.63 | 7.24E-05 | PedA1 |
| 45 | PP_5353 | -1.61 | 1.21E-05 | Tim44-like domain-containing protein |
| 46 | *csiD* | -1.59 | 4.14E-06 | Carbon starvation induced protein |
| 47 | PP_4816 | -1.58 | 3.01E-10 | CidA/LrgA family protein |
| 48 | PP_4379 | -1.58 | 1.56E-13 | 3-oxoacyl-ACP synthase / Beta-ketoacyl-acyl-carrier-protein synthase I |
| 49 | PP_2666 | -1.56 | 1.22E-06 | Rhodanese domain-containing protein |
| 50 | PP_3266 | -1.56 | 3.37E-03 | DUF72 domain-containing protein |

**Table S5: Genes upregulated in *mxtR*-H806N compared to the WT.** Differentially expressed genes with $log_2$foldchange >1 and adjusted *p*-value (padj) <0.05. Gene annotations/functions were retrieved from the Pseudomonas Genome Database (Winsor et al., 2016) and UniProt (UniProt, 2026).

| | Gene / locus tag | log2FoldChange | padj | Annotated function |
|---|---|---|---|---|
| 1 | PP_3843 | 2.47 | 3.73E-21 | Uncharacterized protein |
| 2 | *rplB* | 2.36 | 8.29E-09 | 50S ribosomal protein L2 |
| 3 | *rpsS* | 2.03 | 7.20E-07 | 30S ribosomal protein S19 |
| 4 | *groS* | 1.98 | 8.29E-13 | Co-chaperonin GroES |
| 5 | *rplK* | 1.91 | 2.14E-02 | 50S ribosomal protein L11 |
| 6 | PP_1680 | 1.89 | 1.12E-07 | Alpha-ribazole-5'-phosphate phosphatase |
| 7 | PP_3505 | 1.89 | 6.17E-09 | VWFA domain-containing protein |
| 8 | *groL* | 1.89 | 2.58E-13 | Chaperonin GroEL |
| 9 | *rplV* | 1.87 | 7.70E-06 | 50S ribosomal protein L22 |
| 10 | *cobU* | 1.80 | 1.47E-05 | Nicotinate-nucleotide--dimethylbenzimidazole phosphoribosyltransferase |
| 11 | *cobP* | 1.76 | 1.43E-06 | Adenosylcobinamide kinase / adenosylcobinamide phosphate guanyltransferase |
| 12 | *rpsO* | 1.72 | 9.24E-06 | 30S ribosomal protein S15 |
| 13 | *rplW* | 1.66 | 7.06E-06 | 50S ribosomal protein L23 |
| 14 | PP_3332 | 1.66 | 4.32E-04 | Cytochrome c-type protein |
| 15 | *rplA* | 1.65 | 7.78E-05 | 50S ribosomal protein L1 |
| 16 | *rpsC* | 1.63 | 5.23E-05 | 30S ribosomal protein S3 |
| 17 | *nspC* | 1.62 | 1.51E-05 | Carboxynorspermidine / carboxyspermidine decarboxylase |
| 18 | *rpsF* | 1.62 | 2.91E-06 | 30S ribosomal protein S6 |
| 19 | PP_3784 | 1.62 | 2.24E-04 | Chorismatase FkbO / Hyg5-like N-terminal domain-containing protein |
| 20 | PP_3506 | 1.61 | 3.58E-08 | Magnesium chelatase subunit ChlI |
| 21 | *cobW* | 1.60 | 4.56E-05 | Adenosylcobalmin biosynthesis protein CobW |
| 22 | PP_4978 | 1.58 | 3.01E-15 | Uncharacterized protein |
| 23 | PP_2561 | 1.56 | 5.89E-09 | Extracellular haem-peroxidase |
| 24 | *glcG* | 1.56 | 3.16E-03 | Uncharacterized protein |
| 25 | PP_3235 | 1.53 | 1.71E-04 | CBS domain-containing protein |
| 26 | *rpmJ* | 1.52 | 1.01E-04 | 50S ribosomal protein L36 |
| 27 | *rplJ* | 1.51 | 2.76E-04 | 50S ribosomal protein L10 |
| 28 | *deaD* | 1.50 | 2.70E-04 | ATP-dependent DEAD-box RNA helicase DeaD |
| 29 | PP_2340 | 1.48 | 1.26E-06 | Fe-S protein |
| 30 | PP_3804 | 1.47 | 1.03E-04 | Metal ABC transporter substrate-binding protein |
| 31 | PP_1135 | 1.45 | 2.67E-04 | Uncharacterized protein |
| 32 | *rplD* | 1.44 | 3.26E-04 | 50S ribosomal protein L4 |

| | | | | |
|---|---|---|---|---|
| 33 | *mccA* | 1.44 | 3.90E-10 | Methylcrotonyl-CoA carboxylase biotin-containing subunit alpha |
| 34 | *rpsB* | 1.43 | 3.12E-04 | 30S ribosomal protein S2 |
| 35 | *rpsD* | 1.41 | 1.70E-04 | 30S ribosomal protein S4 |
| 36 | PP_4875 | 1.41 | 4.69E-05 | Membrane protein |
| 37 | PP_1791 | 1.41 | 9.00E-06 | Aldolase/synthase |
| 38 | *rpsJ* | 1.41 | 2.73E-04 | 30S ribosomal protein S10 |
| 39 | *rplU* | 1.39 | 9.92E-06 | 50S ribosomal protein L21 |
| 40 | *ssuD* | 1.39 | 7.10E-10 | Alkanesulfonate monooxygenase |
| 41 | *rpmA* | 1.38 | 3.00E-05 | 50S ribosomal protein L27 |
| 42 | *rpsK* | 1.38 | 5.48E-04 | 30S ribosomal protein S11 |
| 43 | PP_3108 | 1.38 | 2.12E-09 | Rhs-related protein / Tke2 |
| 44 | *atpD* | 1.38 | 9.82E-06 | ATP synthase subunit beta |
| 45 | *cobS* | 1.37 | 2.14E-05 | Adenosylcobinamide-GDP ribazoletransferase |
| 46 | *rpsG* | 1.37 | 3.00E-05 | 30S ribosomal protein S7 |
| 47 | *metP* | 1.36 | 1.80E-06 | L,D-methionine D-methionine ABC transporter permease |
| 48 | PP_5652 | 1.36 | 1.42E-03 | Knr4/Smi1-like domain-containing protein |
| 49 | PP_4858 | 1.36 | 2.23E-03 | Phage infection protein |
| 50 | *rplP* | 1.36 | 4.12E-04 | 50S ribosomal protein L16 |

**Table S6: Genes downregulated in Δ*erdR* compared to the WT.** Differentially expressed genes with $log_2$foldchange <-1 and adjusted *p*-value (padj) <0.05. Gene annotations/functions were retrieved from the Pseudomonas Genome Database (Winsor et al., 2016) and UniProt (UniProt, 2026).

| | **Gene / locus tag** | **log2FoldChange** | **padj** | **Annotated function** |
|---|---|---|---|---|
| 1 | PP_0354 | -6.72 | 3.79E-176 | CBS domain-containing protein |
| 2 | PP_0353 | -6.66 | 3.13E-102 | Exonuclease |
| 3 | *yjcH* | -6.18 | 3.62E-71 | Inner membrane protein |
| 4 | *acsA_I* | -4.62 | 8.55E-150 | Acetyl-CoA synthetase |
| 5 | PP_1635 | -4.48 | 9.58E-76 | Response regulator ErdR |
| 6 | *pqqD_II* | -4.33 | 3.85E-38 | Coenzyme PQQ synthesis protein D |
| 7 | *actP_I* | -3.68 | 1.38E-70 | Acetate permease, Cation/acetate symporter ActP |
| 8 | PP_2861 | -3.40 | 2.81E-46 | Methyl-accepting chemotaxis protein McpP |
| 9 | PP_3179 | -3.28 | 4.65E-36 | LysR family transcriptional regulator |
| 10 | PP_2663 | -3.25 | 3.49E-36 | GfdT protein |
| 11 | *exaE* | -3.22 | 4.03E-25 | PedR2 |
| 12 | *mqo_I* | -3.19 | 2.12E-37 | Malate:quinone oxidoreductase |
| 13 | PP_1587 | -3.04 | 2.28E-18 | NhaP-type Na+(K+)/H+ antiporter |
| 14 | PP_5297 | -2.94 | 8.78E-39 | Amino acid transporter |
| 15 | *scpC* | -2.90 | 1.41E-14 | Acetate:succinate CoA transferase |
| 16 | *sthA* | -2.87 | 4.84E-10 | Soluble pyridine nucleotide transhydrogenase |
| 17 | PP_2671 | -2.71 | 3.34E-16 | Histidine kinase PedS2 |
| 18 | PP_2664 | -2.61 | 1.74E-25 | Sensor histidine kinase |
| 19 | PP_4975 | -2.52 | 1.70E-44 | Long-chain acyl-CoA thioester hydrolase family protein |
| 20 | PP_3014 | -2.48 | 5.17E-28 | Lipoprotein |
| 21 | PP_2666 | -2.30 | 6.80E-07 | Rhodanese domain-containing protein |
| 22 | *aceA* | -2.20 | 1.02E-13 | Isocitrate lyase |
| 23 | PP_2446 | -2.17 | 4.18E-11 | Uncharacterized protein |
| 24 | PP_5659 | -2.10 | 4.19E-20 | Uncharacterized protein |
| 25 | *opdN* | -2.08 | 6.58E-14 | Outer-membrane porin D |
| 26 | *cti* | -1.94 | 1.56E-10 | Esterified fatty acid cis/trans isomerase |
| 27 | PP_2333 | -1.69 | 1.16E-07 | GntR family transcriptional regulator |
| 28 | *yiaY* | -1.64 | 1.89E-12 | Fe-containing alcohol dehydrogenase |
| 29 | PP_2667 | -1.61 | 3.97E-03 | PedC |
| 30 | PP_1832 | -1.56 | 2.98E-05 | Oxidase |
| 31 | *pcaT* | -1.45 | 1.02E-10 | Alpha-ketoglutarate permease |
| 32 | PP_5538 | -1.45 | 6.16E-03 | PedA1 |
| 33 | PP_5365 | -1.38 | 4.84E-10 | Fatty acid methyltransferase |
| 34 | PP_4136 | -1.37 | 3.27E-02 | LuxR family transcriptional regulator |

| | | | | |
|---|---|---|---|---|
| 35 | PP_0333 | -1.34 | 4.10E-02 | Retropepsin-like aspartic endopeptidase RimB domain-containing protein |
| 36 | PP_2669 | -1.34 | 3.56E-03 | PedA2 |
| 37 | *benR* | -1.33 | 1.97E-06 | BenABC operon transcriptional activator |
| 38 | PP_4802 | -1.24 | 2.41E-06 | Lipoate regulatory protein |
| 39 | PP_2588 | -1.22 | 2.35E-03 | Class III aminotransferase |
| 40 | *pycA* | -1.20 | 5.07E-04 | Pyruvate carboxylase subunit A |
| 41 | PP_5620 | -1.19 | 2.10E-02 | MAE-28990/MAE-18760-like HEPN domain-containing protein |
| 42 | *csiD* | -1.14 | 1.62E-02 | Carbon starvation induced protein |
| 43 | PP_0329 | -1.12 | 5.63E-03 | RHS repeat-associated core domain-containing protein |
| 44 | *prpC* | -1.11 | 3.26E-02 | Methylcitrate synthase |
| 45 | PP_0352 | -1.11 | 3.84E-02 | ECF family RNA polymerase sigma-70 factor |
| 46 | PP_4816 | -1.10 | 4.49E-02 | CidA/LrgA family protein |
| 47 | *agmR* | -1.06 | 5.63E-03 | Glycerol metabolism activator |
| 48 | *mmgF* | -1.06 | 4.39E-04 | 2-methylisocitrate lyase |
| 49 | *aldB_II* | -1.03 | 3.82E-02 | Aldehyde dehydrogenase |
| 50 | PP_1083 | -1.02 | 4.56E-02 | (2Fe-2S)-binding protein |

**Table S7: Genes upregulated in Δ*erdR* compared to the WT.** Differentially expressed genes with $log_2$foldchange >1 and adjusted *p*-value (padj) <0.05. Gene annotations/functions were retrieved from the Pseudomonas Genome Database (Winsor et al., 2016) and UniProt (UniProt, 2026).

| | **Gene / locus tag** | **log2FoldChange** | **padj** | **Annotated function** |
|---|---|---|---|---|
| 1 | PP_3843 | 2.58 | 2.12E-20 | Uncharacterized protein |
| 2 | PP_4978 | 1.84 | 5.74E-13 | Uncharacterized protein |
| 3 | *mqo_II* | 1.51 | 1.77E-03 | Malate:quinone oxidoreductase |
| 4 | PP_0899 | 1.24 | 2.67E-06 | Ferredoxin reductase |
| 5 | PP_2561 | 1.23 | 1.45E-03 | Extracellular haem-peroxidase |
| 6 | PP_2323 | 1.02 | 1.41E-03 | N-acetyltransferase domain-containing protein |

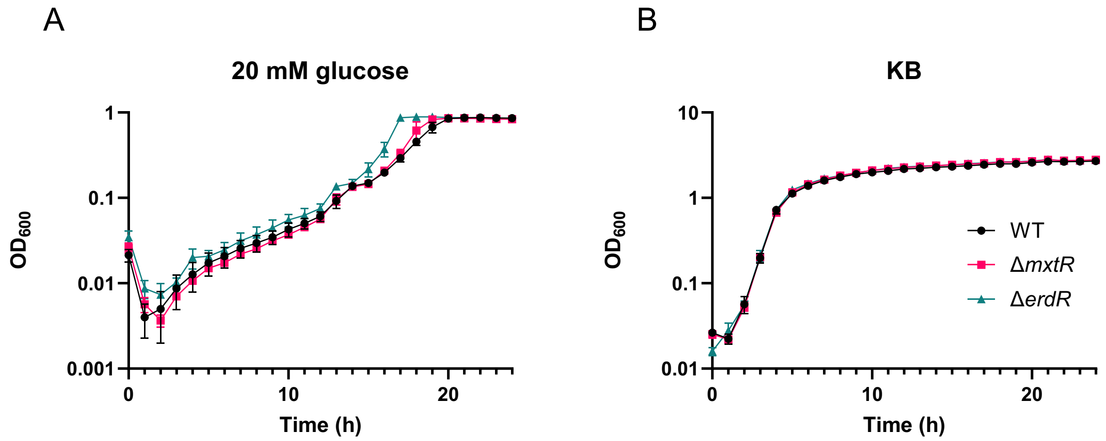


**Fig. S1: Impact of *mxtR* and *erdR* on growth in minimal or complex medium.** (A) Time courses of growth of WT, Δ*mxtR*, Δ*erdR* in M9 minimal medium supplemented with 20 mM glucose or (B) in King's B medium. Growth analyses were performed with an initial $OD_{600}$ of 0.1 and cells were cultivated in 96-well plates in a CLARIOstar Plus plate reader for 24 h, measuring $OD_{600}$ hourly. Mean values and standard deviations of at least three biological replicates are displayed.

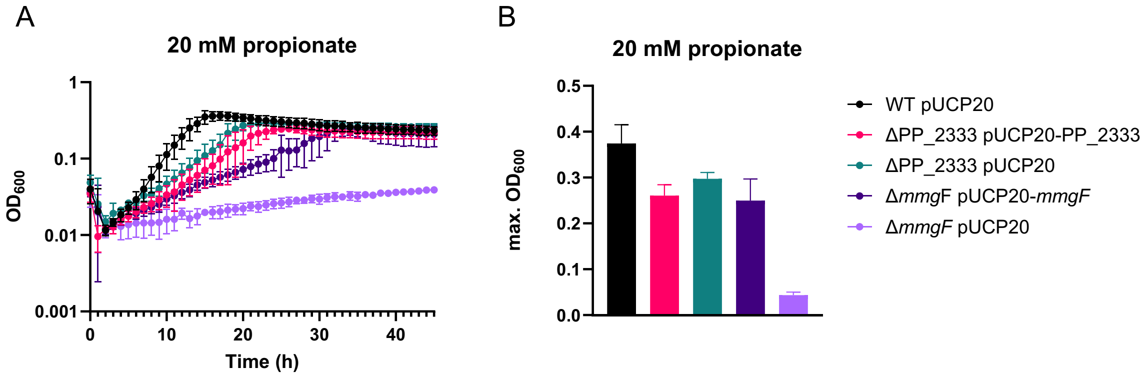


**Fig. S2: Complementation of ΔPP_2333 and *prpE* of the *prp* gene cluster**. (A) Complementation of the growth defect on 20 mM propionate. Growth analyses were performed with an initial $OD_{600}$ of 0.1 and cells were cultivated in 96-well plates in a CLARIOstar Plus plate reader for 45 h, measuring $OD_{600}$ hourly. Complementation of strains was performed with pUCP20-ANT2-PP_2333 or pUCP20-ANT2-*mmgF* and induced with 5 µM anthranilic acid. Strains harboring the empty plasmid pUCP20-ANT2-MCS were used as a control. (B) Maximum $OD_{600}$ reached during complementation growth experiments. Mean values and standard deviations of at least three biological replicates are displayed.

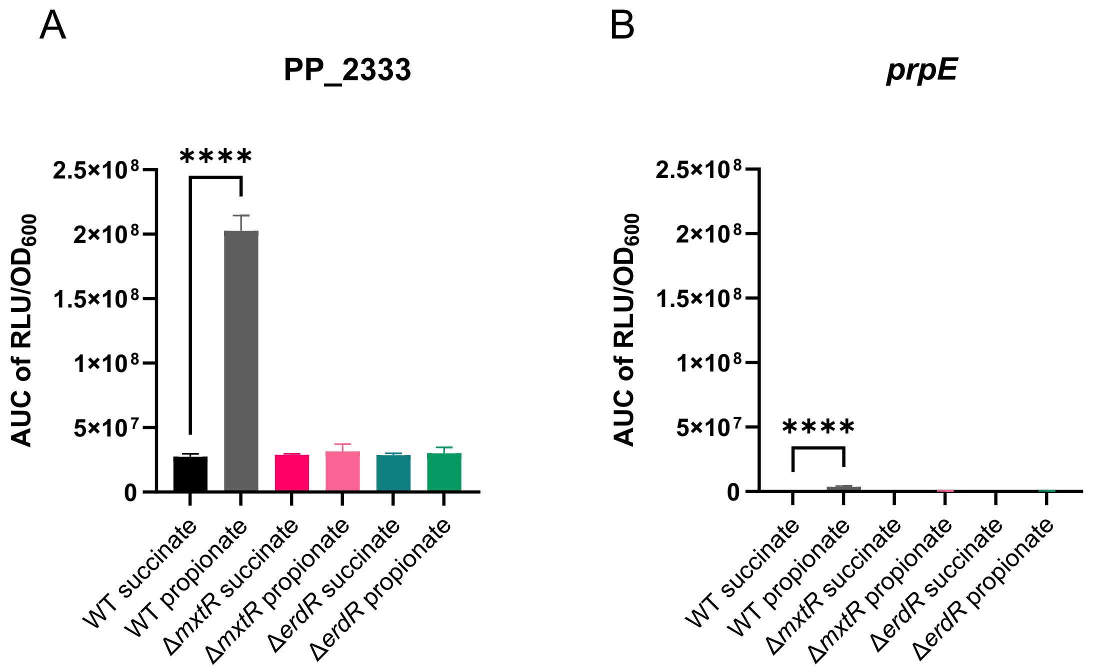


**Fig. S3: Comparison of PP_2333 and *mmgF* expression levels in different carbon sources.** (A) Area under the curve (AUC) of RLU/$OD_{600}$ for PP_2333 or (B) *prpE* (PP_2351) in the WT, Δ*mxtR* and Δ*erdR* strains, cultivated in M9 minimal medium with 20 mM succinate compared to 20 mM propionate. Growth and expression analyses were performed with an initial $OD_{600}$ of 0.1 and cells were cultivated in 96-well plates in a CLARIOstar Plus plate reader for 30 h, measuring $OD_{600}$ quarter-hourly. Mean values and standard deviations of at least three biological replicates are displayed. ****$p$ < 0.0001 (ANOVA and Dunnett's multiple comparisons test).

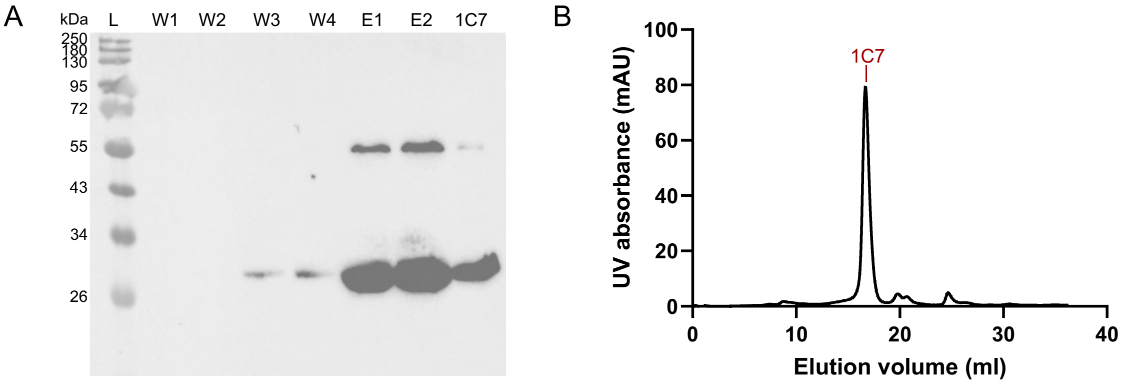


**Fig. S4: Purification of 10H-ErdR via affinity and size exclusion chromatography (SEC).** 10H-ErdR was overexpressed in *E. coli* C41. (A) Western blot of Ni-NTA chromatography wash (W1-W4) and elution fractions (E1-E2), as well as elution fraction 1C7 of SEC. Detection was performed using mouse anti-6xHis primary antibody and chicken anti-mouse IgG HRP secondary antibody. Expected size of 10H-ErdR is 26 kDa. (B) Elution profile of size exclusion chromatography with a Superdex200 column. The fraction with the highest UV absorbance and highest protein concentration is indicated as 1C7. L – protein ladder; W – wash; E – elution fraction of Ni-NTA chromatography. 1C7 – elution fraction of size exclusion chromatography.

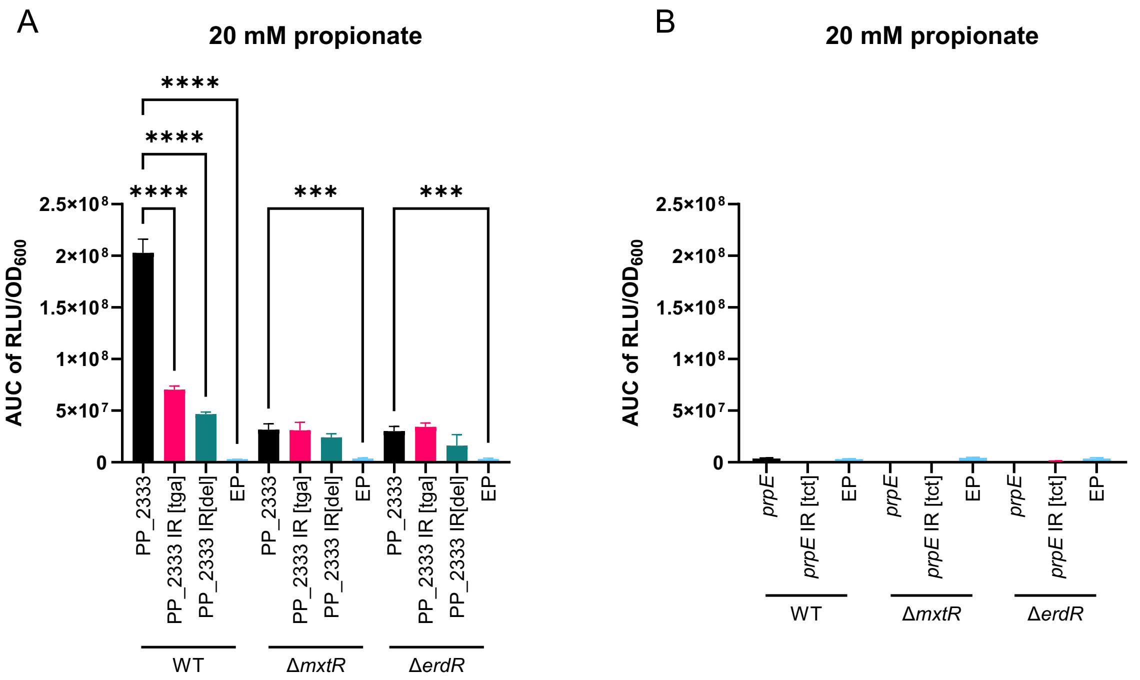


**Fig. S5: Impact of mutation of upstream promoter regions of PP_2333 and *prpE* on expression.** Investigation of modified and unmodified promoter regions of (A) PP_2333 and (B) *prpE* in WT, Δ*mxtR* and Δ*erdR* strains, cultivated in M9 with 20 mM propionate, expressed as area under the curve (AUC) of relative light units per $OD_{600}$ ($RLU/OD_{600}$). Growth and expression analyses were performed with an initial $OD_{600}$ of 0.1 and cells were cultivated in 96-well plates in a CLARIOstar Plus plate reader for 30 h, measuring $OD_{600}$ and luminescence quarter-hourly. PP_2333 – pBBR1-$P_{PP\_2333}$-*lux*; *prpE* – pBBR1-$P_{prpE}$-*lux*; EP – pBBR1-MCS5-*lux*. Mean values and standard deviations of at least three biological replicates are displayed. **** $p < 0.0001$ (ANOVA and Dunnett's multiple comparisons test).

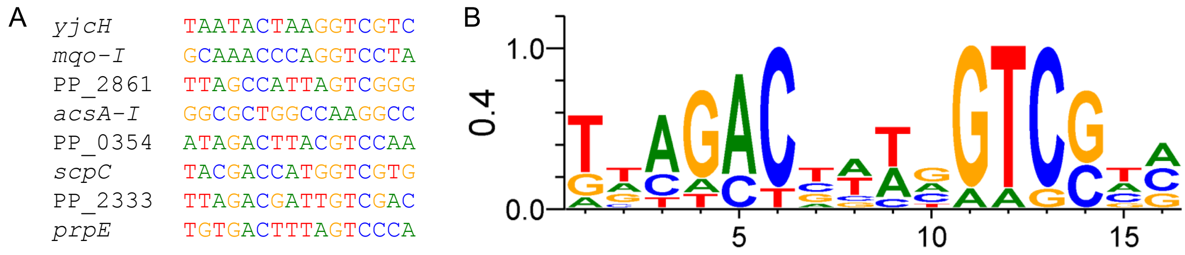


**Fig. S6: Putative DNA binding sites of ErdR of *P. alloputida* KT2440.** (A) Nucleotide sequences upstream of given target genes. The sequences are based on the previously described ACTU motif, a conserved imperfect inverted repeat identified in the putative promoter regions of *acsA* genes of different *Pseudomonas* species (Sepulveda and Lupas, 2017). Disruption of the imperfect palindrome (*prpE*: change to TGT**GAC**TTTA**<u>TCT</u>**CCAT; PP_2333: change to TTA**GAC**GATT**<u>TGA</u>**GACA or deletion of the entire motif) inhibited ErdR binding to the upstream regions and gene expression (Figs. 5 and 6). (B) Sequence logo of the motif, generated using WebLogo 3 (Schneider and Stephens, 1990; Crooks et al., 2004).
